# Emergence of universal computations through neural manifold dynamics

**DOI:** 10.1101/2023.02.21.529079

**Authors:** Joan Gort Vicente

## Abstract

There is growing evidence that many forms of neural computation may be implemented by low-dimensional dynamics unfolding at the population scale. However, neither the connectivity structure nor the general capabilities of these embedded dynamical processes are currently understood. In this work, the two most common formalisms of firing-rate models are evaluated using tools from analysis, topology and nonlinear dynamics in order to provide plausible explanations for these problems. It is shown that low-rank structured connectivity predicts the formation of invariant and globally attracting manifolds in both formalisms, which generalizes existing theories to different neural models. Regarding the dynamics arising in these manifolds, it is proved they are topologically equivalent across the considered formalisms.

It is also stated that under the low-rank hypothesis, dynamics emerging in neural models are universal. These include input-driven systems, which broadens previous findings. It is then explored how low-dimensional orbits can bear the production of continuous sets of muscular trajectories, the implementation of central pattern generators and the storage of memory states. It is also proved these dynamics can robustly simulate any Turing machine over arbitrary bounded memory strings, virtually endowing rate models with the power of universal computation. In addition, it is shown how the low-rank hypothesis predicts the parsimonious correlation structure observed in cortical activity. Finally, it is discussed how this theory could provide a useful tool from which to study neuropsychological phenomena using mathematical methods.

## 1. Introduction

During the last decade, the proliferation of state-of-the-art technologies in the neuroscience community, such as microelectrode arrays, brain-computer interfaces or optogenetics, has given researchers the ability to record, control, and manipulate neural data in unprecedented ways (Cunningham & Yu, 2014; Jazayeri & Afraz, 2017; Vyas et al., 2020). With the previous techniques, it has been possible to obtain huge datasets of neural activity.

A common way to extract useful information from them is to apply dimensionality reduction methods (Altan et al., 2021; Cunningham & Yu, 2014) which in turn allow to recognize that, despite the large dimensionality of the recorded data, neural activation patterns seem to be embedded in a much lower dimensional region, often referred to as *neural manifold* (Gallego et al., 2017, 2018; Sadtler et al., 2014). Indeed, it has been the proceeding of many empirical investigations to obtain neuronal *in vivo* recordings from behaving animals and to subsequentially project the data into this lower dimensional region, in order to infer the most prominent features of the underlying neural dynamics (Chaisangmongkon et al., 2017; Chaudhuri et al., 2019; Churchland et al., 2012; Gallego et al., 2018; Mante et al., 2013).

To understand how neural ensembles perform task-specific computations is the inquiry of much contemporary research, and the role neural manifolds could play in this field is of special int(Cunningham & Yu, 2014; Gallego et al., 2017; Jazayeri & Afraz, 2017). In order to tackle this challenge, many researchers have used recurrent neural networks (RNN) as a model for neural dynamics, searching for a plausible yet tractable modelling capable to shed light on the neural machinery by which cortical computations are biologically implemented (Sussillo, n.d.; van Gerven & Bohte, 2017). These various types of networks describe the behavior of a neural circuit in terms of continuous population variables, primarily the passive somatic potential (Hopfield, 1984), synaptic conductance (B. Ermentrout, 2008; Gort Vicente, 2021), or an average of the firing rate of action potentials (Ermentrout & Terman, 2010a; Wilson & Cowan, 1973), which is why they are commonly referred to as “firing rate models” in theoretical neuroscience (G. B. Ermentrout & Terman, 2010a). Although they are crude approximations of neuronal dynamics that lack many nuances of the mechanisms of spike generation (Gort, 2021), their behavior exhibits many aspects of biological neuronal systems, such as recurrence, feedback, nonlinearity, and principal component activity (Sussillo et al., 2015), and thus they have appealed many neuroscientists who have found in them a way to simulate, interpret, and make sense of empirical data, showing that RNN setups are capable of replicating many experimental observations (Beiran et al., 2023; Chaisangmongkon et al., 2017; Hennequin et al., 2014; Hoellinger et al., 2013; Mante et al., 2013; Meijer et al., 2015; Sussillo et al., 2015; Sussillo & Barak, 2013; Wärnberg & Kumar, 2019))

As far as neural manifolds are concerned, RNN modelling could be a useful framework from which to understand the intrinsic mechanisms that drive neuronal trajectories to settle in a lower dimensional embedding (Jazayeri & Ostojic, 2021).

In support of this proposal, many recent studies have focused on investigating the emergence of neural manifold phenomena in different types of neural networks, some of which having fully structured low-rank connectivity matrices (Beiran et al., 2021, 2023; Dubreuil et al., 2022; Herbert & Ostojic, 2022), while others possessing both a structured low-rank component and an unstructured random weight matrix (Darshan & Rivkind, 2022; Mastrogiuseppe & Ostojic, 2018; Schuessler et al., 2020).

Although interesting progress has been made in this direction, the approaches used so far rely mainly on computational tools (Maheswaranathan et al., 2019) and statistical methods, such as mean-field theory (Beiran et al., 2021; Mastrogiuseppe & Ostojic, 2018; Schuessler et al., 2020). In some respects, there exist limitations in establishing sufficient conditions for the existence of neural manifolds and measuring their dimensionality (Altan et al., 2021). For example, the mean-field theory approach works only in the limiting case when the number of neurons tends to infinity (Sompolinsky et al., 1988), and thus cannot fully determine the evolution of a given finite-size neural system. Similarly, computational perspectives cannot make general statements concerning the mechanisms that unfold at the level of a large class of neural models, as they can only provide concrete numerical insights into a given system. Therefore, in these paradigms it is difficult to obtain general results, and one has to be content with interpreting the implementation of specific computational tasks in terms of concrete model configurations (Sussillo & Barak, 2013), without being able to make general statements about the universal computational capabilities with which neural networks are endowed.

Moreover, due to these pitfalls, most of the aforementioned works had to limit the size of the studied neural manifolds to as few as 2 variables (Herbert & Ostojic, 2022; Mastrogiuseppe & Ostojic, 2018; Schuessler et al., 2020), while empirical investigations show that these subspaces can consist of up to 10 independent components (Sadtler et al., 2014).

In the present study, we will consider the two best known formalisms of firing-rate models to study in detail their behavior from the point of view of dynamical systems theory, topology and analysis. We will prove that, for fully structured low-rank connectivity matrices, both models exhibit invariant and globally attracting manifolds whose intrinsic dimensionality corresponds to the rank of the connectivity matrix.

We will then study the computational performance that results from low-rank models and show that they are universal approximators of input-driven control dynamical systems, extending results that have so far focused mainly on the autonomous case.

This will lead to announce some general results on the mechanisms underlying some important internal processes, such as the storage, control and production of muscle activity; the generation of attractive memory states and central patterns; and the universality to simulate effective procedures using serial symbolic manipulation with finite memory capacity. Finally, it is going to be shown how the low-rank connectivity hypothesis leads to empirically contrasted predictions, and the significance of these results is going to be discussed from a cognitive perspective.

## 2. Results

In this section, we will present statements concerning the dynamics of the following mathematical models:

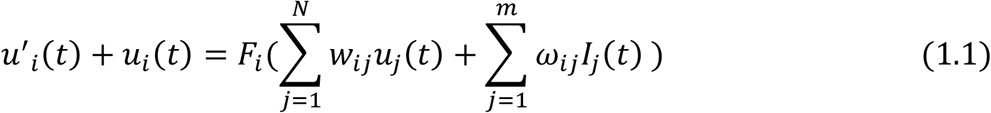

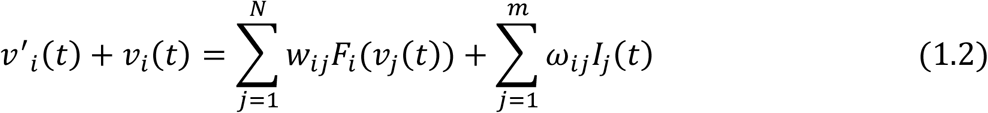

The first one describes the time evolution of a filtered measure of the firing rate (Miller & Fumarola, 2012), while the second does so for the voltage of the cell when action potentials are removed (Ermentrout & Terman, 2010a). Each *F*_*i*_ models the relationship between the voltage and the spike frequency of each neuron, and they are going to be called transfer functions. The parameters, *w*_*ij*_, *ω*_*ij*_ stand for the strengths of the synaptic projections coming from circuit and input neurons, respectively. We will often refer to the first formalism as the u-model or of the Wilson-Cowan type, and the second as the v-model or of the Hopfield type.

For the sake of simplicity, the above equations assume that all the nodes of the net have equal time constants. We will further suppose that for every *F*_*i*_, there exists a scalar, *θ*_*i*_, which stands for each neuron’s action potential threshold, such that *F*_*i*_(*x*) = *σ*(*x* − *θ*_*i*_), being *σ*: ℝ → (0,1) a sigmoid function. The following results, however, hold for more general transfer functions as well.

The above formalisms are not the only ones that attempt to model collective neural behavior using physiologically relevant, continuous neural variables. However, they are analytically tractable and can be found among the most widely used models in systems neuroscience. Similar but more biophysically realistic models, also based on continuous variables and neural population measurements, can be found in (Pasquale, Sussillo, Abbott, 2023; Ekeberg, 1993; Thalmeier et al., 2016).

### 2.1 Conditions sufficing the formation of neural manifolds

In many cases, both computationally and experimentally, it is common to visualize data through dimensionality reduction methods (Hennequin et al., 2014; Sussillo & Barak, 2013). Although their estimates are good enough for many practical reasons, they can sometimes be deceptive about the geometry or dimensionality of the studied lower-dimensional structures, especially when linear methods are involved (Altan et al., 2021).

For instance, even in the case of a network exhibiting chaotic high-dimensional fluctuations, which is known to be the dominant scenario for large recurrent networks with unstructured random connectivity (Mastrogiuseppe & Ostojic, 2018; Sompolinsky et al., 1988; Stern et al., 2014; Williamson et al., 2016), the solutions could settle on a strange attractor forming a relatively correlated data set that would subsequently yield some eigenvalues close to zero when performing a principal component analysis (PCA). However, this would not entail the dynamics to remain in a low-dimensional subspace, since the attractor itself could be full-dimensional, as numerical evidence suggests in figure 2.

**Figure 1.**
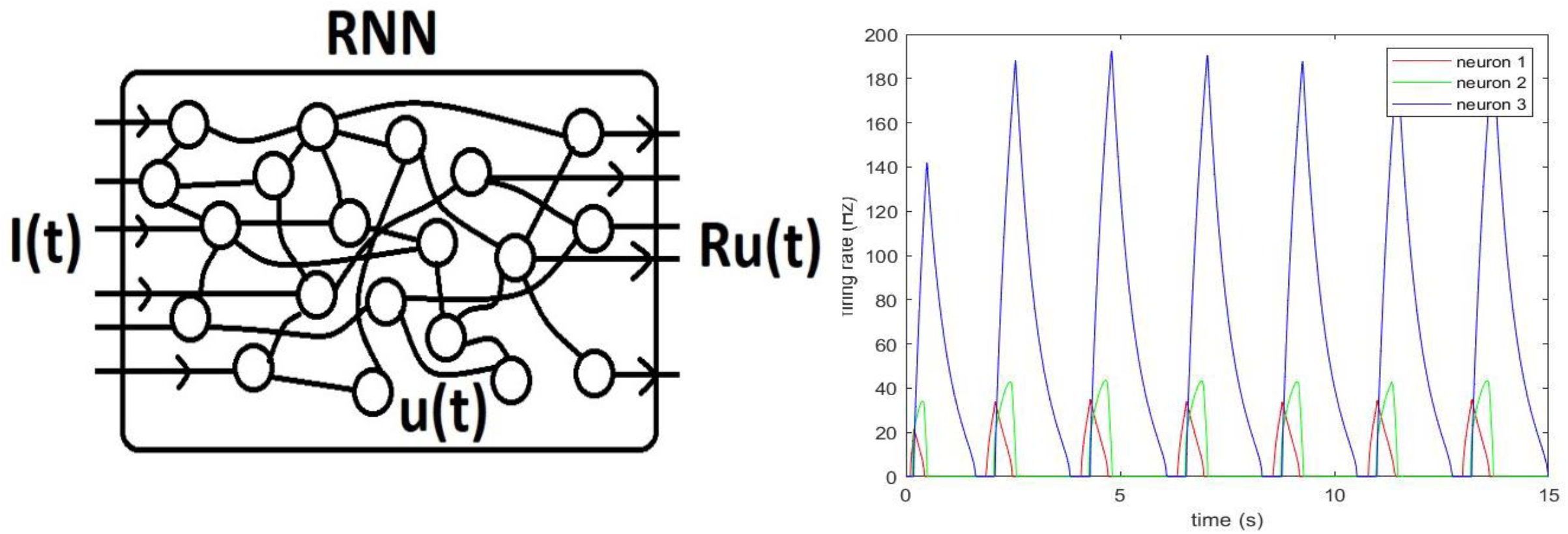
Left: a schematic representation of a recurrent neural network, being I(t) the input vector; u(t) the internal state array, formed by the measures of the activations of the recurrently connected nodes; Ru(t) the readout of the circuit, which extracts the output of the system through the action of a linear operator, R. Right: Plot showing the firing rate of a three-neuron network wired to simulate the periodic bursting of the Tritonia Diomedea central pattern generator. The model shows an intrinsic oscillator such as the one in the original circuit, which equips the organism with an escape swimming response. See (Katz, 2009) for information about the biological circuit, and [16] for details on the computational modelling.

**Figure 2 :**
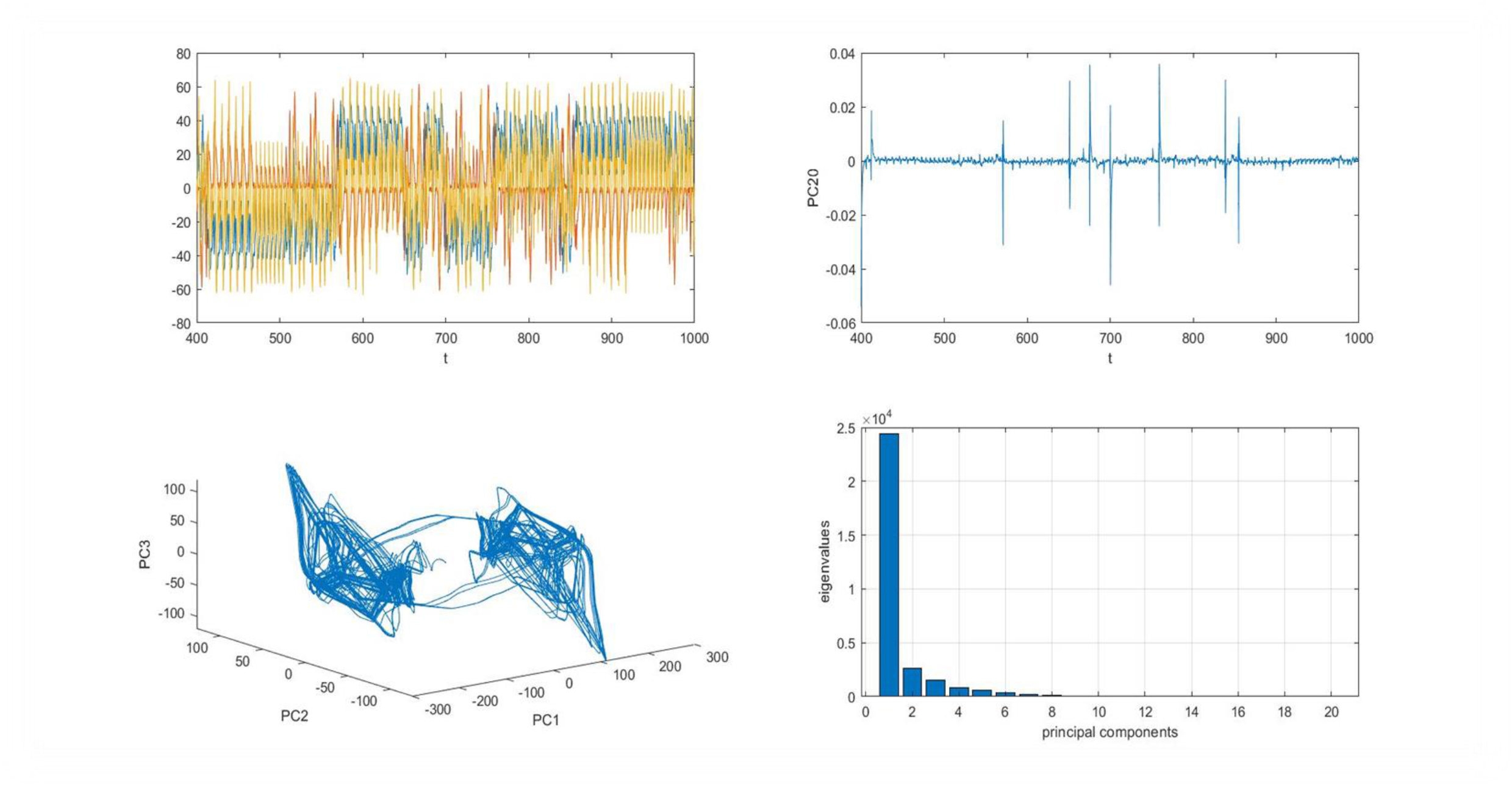
Simulation of a 20 neuron Hopfield network whose connectivity was randomly wired for a given initial condition. In the first column, it is shown the activation of 3 units displaying chaotic behavior, and below the underlying double-scroll attractor is projected onto the 3 first principal components. In the second column, the evolution of the last principal component is plotted together with a histogram showing the eigenvalues of the covariance matrix. Although the last 10 eigenvectors could be neglected in a lower-dimensional model of the dynamics which could explain almost the whole variance of the system, the last component is nonconstant, and thus the network exhibits full-dimensional chaotic fluctuations.

**Figure 3:**
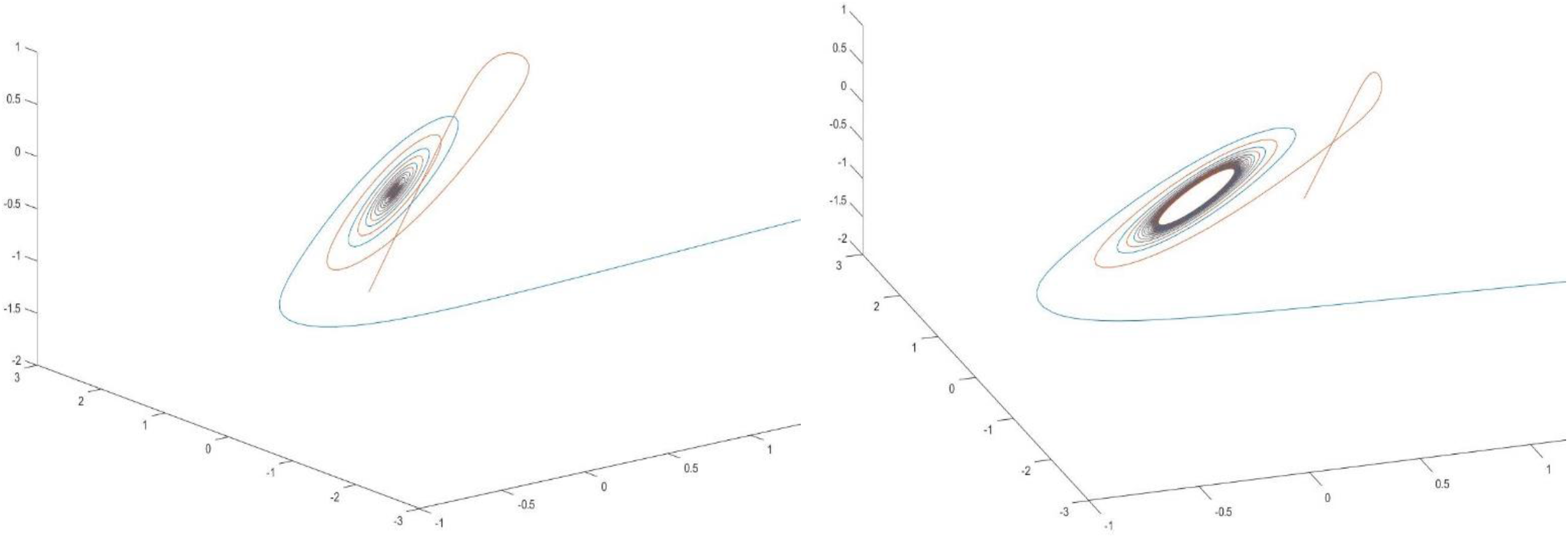
Two trajectories of a 3 neuron Wilson-Cowan system undergoing a Hopf bifurcation on a 2-dimensional manifold. The bifurcation parameter is a single weight whose modification leaves rank(W)=2 unchanged, so that the emergence of a globally attractive surface can be observed in both cases, according with the conclusions of theorem 1.1.

**Figure 4:**
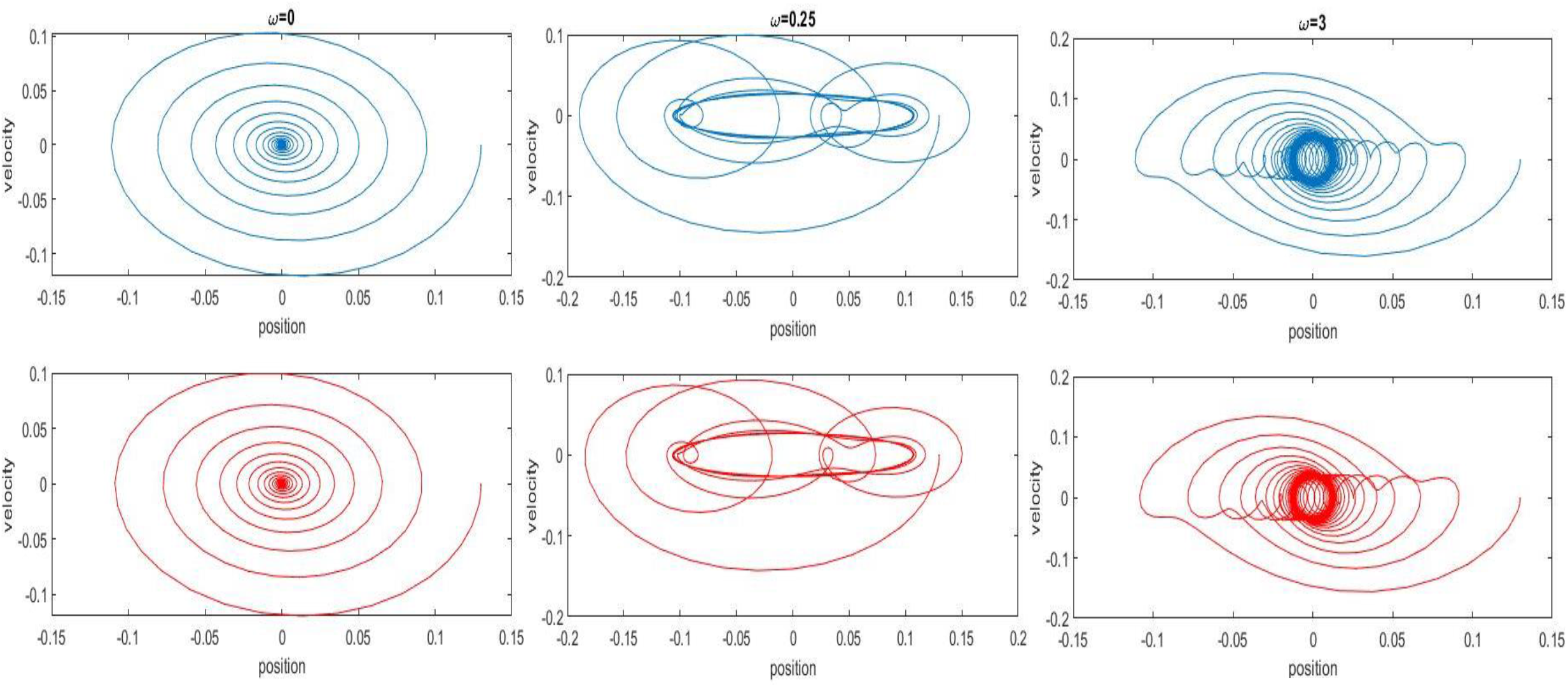
projection of a phase plane trajectory of a periodically driven damped harmonic oscillator. In the first row, in blue, the solutions of the original system; in the second row, in red, an approximation performed with a firing-rate network of the Wilson-Cowan type. Each column represents a different frequency for the driving force.

**Figure 5:**
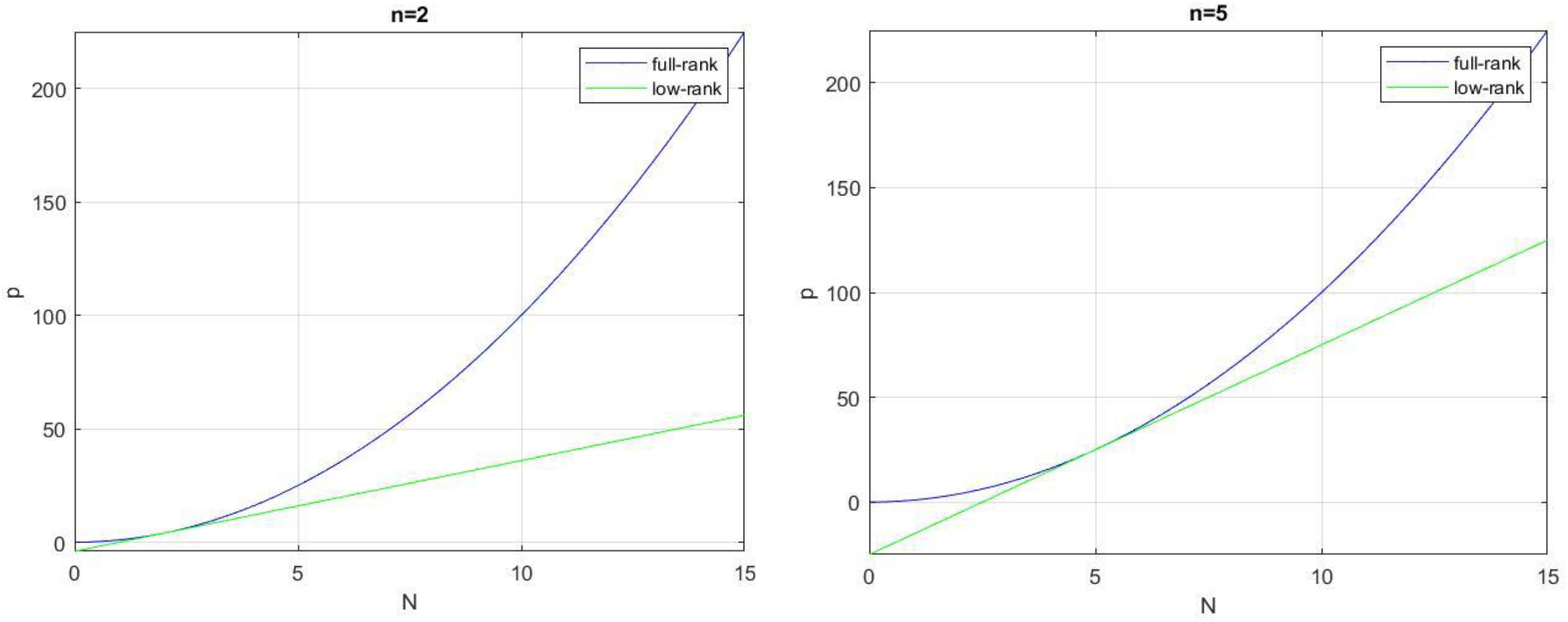
plots showing the number of parameters, p, needed to fully specify a connection matrix, versus the number of neurons, N, of the network. Different dimensionalities for the simulated dynamical systems, n, are supposed. It can be seen how the number of parameters needed in the low-rank setup is of order N, whereas in the full-rank scenario p equals the number of matrix components, N^2^. It is further shown in the figure, as it can be readily seen in the derived expressions for p, that the low-rank function for p corresponds to the tangent line of the full-rank curve, whose tangency point is given by the number of dimensions of the implemented dynamics.

It is thus that we are interested in seeking conditions accurately predicting the formation of neural manifolds. As previously announced, this will be achieved by restricting weight matrices to be non-invertible. Thereby, we present the following statements:

#### Theorem 1.1

Let *W* be the connection matrix of a given u-model with constant input. Then, it possesses an invariant, globally attractive manifold, ℳ, which can be parametrized by a homeomorphism *f*: ℝ^*rank*(*W*)^ → ℳ.

The proof of this theorem relies on constructing a Cauchy sequence of functions converging to the map *f*.

#### Theorem 1.2

Let *W* be the connection matrix of a given v-model with constant input. Then, it possesses an invariant, globally attractive manifold whose intrinsic dimension is equal to *rank(W)*. Furthermore, this manifold is Euclidean.

For the proofs, see methods, 5.1. Something that can be inferred from the previous theorems is that, although the manifolds emerging from each model could be geometrically different, they are always homoeomorphic. This does not imply that all observed neural manifolds must be topologically Euclidian, as on the contrary, for example, in the head direction circuits of mammals and insects ring attractors have been observed (Chaudhuri et al., 2019; Kim et al., 2017). Nevertheless, these topologically more complex structures could well be considered as sub-manifolds embedded in the manifolds described in the previous theorems. Section 2.2 is going to study the dynamics evolving in these embeddings, showing that the submanifolds they enclose can be very rich indeed.

From now on, we will reserve the term “neural manifold” for the topological Euclidean manifolds described in the previous theorems, and we will refer to the dynamical systems evolving on these invariant regions as neural manifold dynamics.

So far, we have studied u-models and v-models separately. Now that explicit conditions predicting the existence of dynamical embedding manifolds have been found, it could be asked what relation exists, if any, between neural manifold dynamics in both kinds of models. The following proposition addresses this problem:

#### Proposition 1.1

Suppose we have a u-model and a v-model both with equal transfer functions, connectivity arrays and constant inputs. Then, their neural manifold dynamics are topologically conjugate.

We say that the set of solutions of two systems are topologically conjugate if there is an invertible bicontinuous transformation that yield a trajectory of one system whenever applied to a solution of the other one. More technically, it means that there exists a homeomorphism which commutes with the flow map. See methods, section 5.1 for the proper definition, or (Hirsch et al., 2013a).

Intuitively, two dynamical systems being conjugate indicate that they are topologically equivalent, meaning their qualitative behavior remains unchanged, so that things like the number of fixed points, their stability, the presence of cycles or connection orbits as well as the existence of chaotic regions in state space are preserved by moving between the systems. This result provides a useful analytical tool for investigating the behavior of one type of network knowing the behavior of the other.

For example, the well-established fact that randomly wired networks of Hopfield neurons tend to behave chaotically as their dimensionality increases (Sompolinsky et al., 1988) can now be easily proved for Wilson-Cowan networks - we only have to use the fact that the most commonly used definitions of chaos persist under conjugacy (Lu et al., 2013) and apply proposition 1.1.

Ultimately, this theorem assures that u-models and v-models are mutually consistent, being both equivalent expressions of the same process, from a topological perspective. In Miller & Fumarola, 2012, it was proven that both models were related via an affine map, but it could not be provided a way to find a complete trajectory of the u-model given a solution on the neural manifold of the v-model. In the previous proposition, it was proven that such an invertible connection between the networks exists in all cases, broadening the previous work whenever constant inputs are considered. See methods, section 5.1, for the proof and information about the conjugacy.

### 2.2 Exploring the universality of low-dimensional computations

The fact that low-rank structured systems have invariant manifolds implies that a dynamical system evolves within these hypersurfaces. In this section, we will explore the nature of these emergent, populationally distributed processes and the extent to which they can endow cortical systems with useful computational mechanisms.

From a dynamical systems perspective, it would be interesting to characterize both geometrically and topologically the various phenomena that take place in these manifolds. With respect to this problem, it is found, surprisingly, that neural manifold dynamics are, in some sense, dense in the space of dynamical systems, which means that for each system of differential equations there is a neural ensemble whose projected dynamics approximates the system up to a given, yet arbitrarily small, degree of precision. This is stated more accurately in the following theorems, one for each of the models studied:

#### Theorem 2.1

Let Ω ⊂ ℝ^*n*^ be compact, *G* ∈ *C*^1^(ℝ^*n*^ × ℝ^*m*^, ℝ^*n*^) and *x*(*t*) the solution to the initial value problem

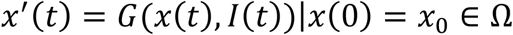

being *I*: ℝ → ℝ^*m*^ drawn from an uniformly bounded set of continuous functions. Then, there exists *N* ∈ ℕ, a matrix *R* ∈ ℝ^*n*×*N*^ and a u-model with input *I*(*t*) and connectivity *W* such that, for any *ε* > 0 and any *T* > 0, being [0, *T*] included in the maximal interval of existence, it is fulfilled that

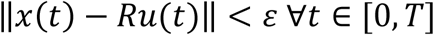

given appropriate initial conditions, *u*(0). Furthermore, *rank*(*W*) = *n*.

#### Theorem 2.2

Let Ω ⊂ ℝ^*n*^ be compact, *G* ∈ *C*^1^(ℝ^*n*^ × ℝ^*m*^, ℝ^*n*^) and *x*(*t*) the solution to the initial value problem

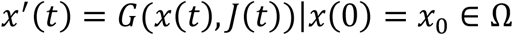

being *J*: ℝ → ℝ^*m*^ drawn from an uniformly bounded set of differentiable functions. Then, there exists *N* ∈ ℕ, a matrix *R* ∈ ℝ^*n*×*N*^ and a v-model with input *I*(*t*) = *J*(*t*) + *J*^′^(*t*) and connectivity *W* such that, for any *ε* > 0 and any *T* > 0, being [0, *T*] included in the maximal interval of existence, it is fulfilled that

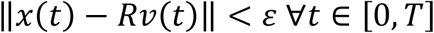

given appropriate initial conditions, *v*(0). Furthermore, *rank*(*W*) = *n*.

The proofs of these theorems rely on the universal approximation theorem from artificial neural network theory.

In the foregoing, the matrix R represents the readout which defines the output of the network once it is applied to the vector of neural states. The fact that rank(W)=n predicts, using the results of section 2.1, the emergence of stable invariant manifolds in the systems studied, at least for the case when the emergent dynamics are autonomous. This establishes a link between the theory of neural manifolds and the universal approximation properties of firing rate models, which will be further explored in the following subsections.

The results we have just outlined can be summarized saying that, for any driven nonautonomous dynamical system, there exists a neural network of each type whose outputs can approximate its solutions with arbitrary accuracy and during arbitrarily long time periods. This is a property that provides our neural models with a great power of simulation and control. Biological neural networks must be able to perform a variety of different tasks, as they alone are responsible for a wide range of internal processes, ranging from the generation and control of the muscular patterns that constitute observable behavior to the computations that mediate between the stimulus and responses. For this reason, we will now turn to the behavioral and cognitive consequences that follow from the results obtained so far.

Rigorous propositions with relevant biological interpretations will be presented, concerning the coordination and execution of muscle patterns, the storage of mnemonic and behaviorally relevant attracting states and the universal implementation of symbolic procedures.

#### 2.2.1 Storing compact sets of muscular trajectories

The generation of muscular activity patterns is one of the most studied topics within the dynamic systems approach to large-scale neuroscience (Vyas et al., 2020). This perspective has provided experimental and computational evidence suggesting that motor areas store large sets of muscle patterns whose production is triggered by switching between initial conditions during preparatory activity (Churchland et al., 2012; Hennequin et al., 2014; Michaels et al., 2016; Sussillo et al., 2015). It is known that there is a strong functional correlation between muscle activity and the primary motor area of the cortex (Gallego et al., 2018; Miri et al., 2017) so that the dynamic properties of the latter can be readily mapped onto the temporal development of the former.

Here we will use the aforementioned theorems to make a statement about the large-scale storage of muscle patterns. Each of these responses is going to be modelled as a trajectory represented by a continuous mapping of the form *r*: [0, *T*] → ℝ^*n*^, *T* > 0. The set of all these trajectories are going to be equipped with the uniform norm:

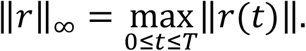

##### Proposition 2.1

Given any compact set of functions under the uniform norm, 𝒜, whose elements are trajectories of the form *r*: [0, *T*] → ℝ^*n*^, *T* > 0, and given any *ε* > 0, there exists a u-network (respectively a v-network) such that, for every *r* ∈ 𝒜, its output, given appropriate initial conditions, fulfils that

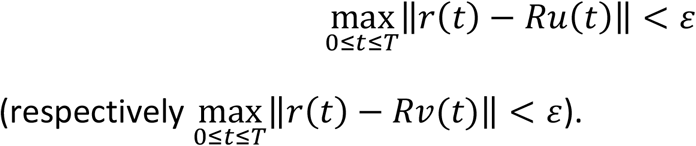

This proposition claims that it is possible to develop a neural system capable of implementing arbitrary compact sets of muscle patterns, the production of which is preceded by pinpointing an appropriate initial condition, as claimed in previously mentioned studies (Churchland et al., 2012; Hennequin et al., 2014; Michaels et al., 2016; Sussillo et al., 2015).

Our model of muscle activity is essentially autonomous and is controlled by the assignment of initial conditions. Therefore, theorems 1.1 and 1.2 predict that the manifold in which the relevant muscle pathways are embedded should be preserved and have the same geometry regardless of the evoked pattern or the elapsed time. This has indeed been reported by experimental studies (Gallego et al., 2018).

Moreover, the fact that autonomous and continuous dynamics evolving along neural manifolds follow the conditions of Picard’s existence and uniqueness theorem implies that two different trajectories should never cross each other. This imposes conditions on the intrinsic dimensions of the manifold, which should be provided with additional degrees of freedom in order to disentangle different trajectories, as shown in methods, section 5.3. The existence of such control directions has also been reported earlier in empirical investigations (Hall et al., 2014; Russo et al., 2018).

Taken together, we believe that these results provide some initial evidence supporting our hypothesis.

#### 2.2.2 Attracting sets of neural activity

One limitation of both theorems 2.1 and 2.2 is that, although the approximation of flows is universal and arbitrarily accurate, it can only be so, in general, for a time interval of limited duration. Nevertheless, this can be overcomed by providing stronger hypothesis. This will be achieved by assuring that all the approximated solutions eventually converge to stable limiting trajectories, as the following result suggests:

##### Proposition 2.2

Suppose we have an autonomous system of differential equations induced by a function *G* ∈ *C*^1^(ℝ^*n*^, ℝ^*n*^). Let *U* ⊆ ℝ^*n*^ be open and Ω ⊂ *U* be a compact domain, and suppose further that there exist a family of *l* ∈ ℕ periodic orbits, 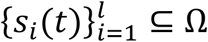, such that, if *x*(*t*) is a solution given an initial condition *x*_0_, then

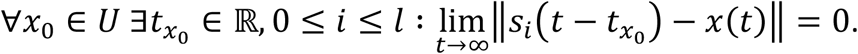

In this case, given *ε* > 0 and any trajectory, *x*(*t*) with *x*(0) ∈ Ω, there exists a u-network (respectively a v-network) whose output, given appropriate initial conditions, fulfils that:

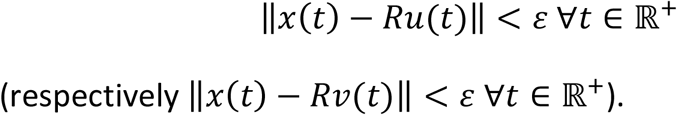

This allows to extend the previous results for all forward time, allowing our neural models to approximate phase trajectories *ad infinitum* whenever the target system is endowed with stable periodic orbits, being these either fixed points or limit cycles.

Attracting fixed points are traditionally used in the field of artificial neural networks as a tool for implementing auto-associative memory devices (Hopfield, 1982) which curiously often resort to low-rank weight matrices, just as in our approach (Amit et al., 1985; G. B. Ermentrout & Terman, 2010b), although there the matrices are further restricted to be symmetric, unlike in our case. This splitting of the neural state space into different basins of attraction has been observed in the study of context-dependent episodic memory in the hippocampus (Wills et al., 2009).

Attracting fixed states of persistent neural responses also underpin much contemporary research on working memory, where stable bumps of neural activity have been found in the prefrontal cortex of animals prior to retrieval of task-related stored information (Wimmer et al., 2014).

Beyond fixed points, attractive limit cycles are also important objects in neurodynamics, as they are responsible for many robust oscillatory phenomena, such as in locomotion or respiratory patterns. An important class of biological neural circuits that have been shown to sustain intrinsic limit cycle oscillations are the so called central pattern generators (CPGs), which spontaneously support endogenous and repetitive fluctuations without the need of external periodic driving (Yuste et al., 2005).

These different types of neural phenomena all have in common that they are based on locally attracting periodic phase trajectories, as the ones assumed in the hypothesis of proposition 2.2.

Therefore, this result encompasses all these phenomena in the context of the universal approximative capabilities of firing rate networks.

The statement of the previous proposition does not only intend to show the ability of connectionist models to implement attracting memory states or CPGs, which has already been achieved (Hoellinger et al., 2013; Hopfield, 1984; Hopfield & Tank, 1986; Ijspeert, 2001), but also to demonstrate their universality in accomplishing these tasks, to prove that they can behave in this way persistently, and to show that an effective way to implement these functions is to use low-rank connectivity patterns, from which neural systems are expected to uphold neural manifold dynamics, as it has already been observed experimentally.

#### 2.2.3. Universal simulation of finite memory Turing machines

In the study of computable functions, i.e., mappings for which there is an effective procedure to achieve the corresponding outputs, classical computational theory has relied on the Church-Turing thesis, which basically states that any algorithm can be implemented by a Turing machine, and thus relies on this theoretical construct in order to tackle many theoretical problems in computational science.

Intuitively, a Turing machine is composed of an infinite tape consisting of many ordered symbols borrowed from a finite set, the alphabet, where only one of them, the empty symbol, is allowed to occur infinitely often; a set of states, one of which is the initial and some others the final acceptors; and a hypothetical device that can move along the tape so that once it has read a symbol, it is able to rewrite the current cell of the tape and move in either direction to a new cell once it has updated its state, thus implementing a transition function. For a deeper understanding of these concepts we refer the reader to any textbook about computation theory, such as (Lewis & Papadimitriou, 1998).

The tape of the Turing machine store the problem’s information given some formal language (Lewis & Papadimitriou, 1998), which the moving lecture device has the job to overwrite so as to terminate the computation. This sequentially manipulated tape, once the alphabet elements are indexed, can be thought as a natural number. We will say that a Turing machine have finite memory if it only performs interpretable computations on some compact set of ℕ, this is, if it is only defined over some set of tapes whose non-blank symbols are pushed to some common initial interval of finite length.

It is the quest of this subsection to explore firing rate networks power for serial symbolic computation. Much of this effort will rely on (Branicky, 1995), where it was proven that continuous time dynamical systems on ℝ^3^ have the power of universal computation, meaning they can simulate any arbitrary Turing machine. The way such simulation can be set up is by encoding each Turing machine’s configuration (this is, the present state, the whole infinite tape and the current cell location) as an open domain in phase space, and then to define the flow of the system such that it emulates the same transition function as that of the original Turing machine. See (Branicky, 1995) or section 5.3 for more details. Mixing this idea with our previous results give rise to the following statement.

##### Proposition 2.3

Given any compact set *S* ⊂ ℕ and any computable function *f*: *S* → ℕ, there exists a u-model (respectively a v-model) capable to implement an effective procedure for *f*. This simulation lies on a 3-dimensional, invariant and globally attractive manifold.

Using the Church-Turing thesis, we can think of any effective procedure as the operations performed by a Turing machine. Following the previous notions of simulation, we can think of the settings of this Turing machine as open regions in the predicted 3-D neural manifold, being those sets of states sewed together by the phase trajectories implementing the transition function. These trajectories would be in charge to transit between different domains of the state space with distinct symbolic interpretations. Interestingly, tunnelling trajectories joining different state space regions have been showed to implement robust computations in recordings of the monkey cortex during decision-making tasks (Chaisangmongkon et al., 2017).

Also backing what is been established experimentally, this proposition shows that it is not necessary to have great dimensionality in order to implement powerful neural computations (Gallego et al., 2017; Mante et al., 2013; Sadtler et al., 2014).

The fact that we have to settle for finite-memory computation is due to the fact that Turing universality is a theoretical notion, since no physical system such as neural circuits or modern computers can store infinite memory. Indeed, theoretical papers proving Turing universality of traditional discrete time neural networks either assume that these networks have unbounded precision (Sontag & Siegelmann, 1991), which is physically implausible, or that they can recruit more neurons, indefinitely, whenever they need more memory (Chung & Siegelmann, 2021), which make the size of these networks effectively infinite.

In the proof, section 5.3, it can be seen that our construction is robust under small perturbations of the neural trajectories, so that it could be, hypothetically, implemented by real noisy neural systems with intrinsic bounded precision.

### 2.3. Implications of low-rank connectivity

The fully structured low-rank hypothesis that it was imposed on the connectivity as to show both the existence of neural manifolds and the emergence of computationally useful dynamics constitutes a sufficient condition for these purposes, but not a necessary one. Indeed, since invertible matrices form an open and dense subset in ℝ^*N*×*N*^, one could arbitrarily approximate any non-invertible connectivity array by a full-rank matrix, therefore obtaining some full-rank connectivity matrix with the same approximation power of the non-invertible one, thus discarding converse results. Nonetheless, the restriction of low-rank wirings could well have positive benefits for neural ensembles over other kinds of setups (Beiran et al., 2023).

For instance, take the number of parameters, say *p*, needed in order to fully define the connectivity matrix when implementing an *n* dimensional dynamical system. This measure will obviously depend on the size of the network, this is, the total number of neurons, *N*. If we consider that connectivity could lie anywhere in the *N* × *N* matrices space, the number of independent parameters needed to define the connections would be given by *p* = *N*^2^, regardless of the dynamical system’s dimensionality. However, if we just take the family of *n*-rank matrices into consideration, as theorems 2.1 and 2.2 suggest, since we just need to define *n* columns and the rest of them turn out to be linear combinations of the former, the amount of independent parameters needed in order to fully define the connections, this time, works to be given by *p* = *n*(2*N* − *n*).

Therefore, the number of degrees of freedom of the matrix depend linearly on *N*, instead of doing so quadratically, thus regularizing the connectivity in a way it can implement difficult tasks relying on a structured parsimonious connectivity of reduced complexity. Indeed, it’s been shown that low-rank structured networks generalize their behavior to novel stimulus better than their full dimensional counterpart (Beiran et al., 2023).

An appealing characteristic of RNN modelling is that it allows both to explain specific aspects of experimental data and to generate empirically testable predictions (Jazayeri & Ostojic, 2021; Vyas et al., 2020; Wärnberg & Kumar, 2019). These kinds of explanations have been emphasized in this study, where hypothetical mechanisms for the implementation of muscular control, attracting memory states, central pattern generation and serial symbolic processing were developed.

On this matter, interesting predictions can be done based on the low-rank regularization hypothesis. For instance, suppose we would like to know the value of some descriptive statistics of the model, like the correlation between pairs of individual units. This could be done individually for each solution, subsequently averaging over some set of initial conditions in neural state space (see methods, 5.4). Under the low rank hypothesis, and given the existence and uniqueness theorem, any of these solutions would be completely specified once determined the initial conditions plus the *p* = *n*(2*N* − *n*) number of parameters needed to fully determine the underlying connectivity. Even if we included individual firing thresholds and membrane time constants to the model, once we average over the relevant set of initial conditions the number of independent parameters needed in order to fully determine the correlation matrix of the model will be of order *N*, instead of the order *N*^2^ values one would expect in the full-rank structure and in the random connectivity cases.

It turns out this somewhat surprising result have been empirically confirmed in the upper sensory cortex, where it was shown that the *N* correlations from individual neurons to the summated activity of the whole network suffice to determine a wide fraction of the entire matrix of pairwise correlations between network cells (Okun et al., 2015), showing it is only needed order *N* independent parameters so as to grasp the whole order *N*^2^ dimensional correlation structure.

Again, here the low-rank hypothesis is a sufficient condition, but not a necessary one. Indeed, an alternative stronger hypothesis would be to suppose neural data remained on a completely flat *n-*dimensional surface, as then a straightforward PCA would show the covariance matrix, and thus also that of the correlations, would have rank equal to *n*, and therefore the number of independent defining parameters would also be of order *N*, using preceding arguments (methods 5.4).

Nonetheless, restricting the neural manifold to be flat is a rather strong and unrealistic assumption, as even in the v-model the observed manifold is not Euclidian, since to obtain the firing rates we still must apply a non-linear transfer function, *F* (Ermentrout & Terman, 2010a). Thus, the low-rank hypothesis turns out to be a plausible frame from which to make sense of the seemingly parsimonious correlations observed between individual neurons in the cortex. Indeed, in (Okun et al., 2015) the connectivity matrix was already assumed to be low-rank, since in order to explain how enhanced nonspecific connectivity increases population coupling it was assumed that inhibitory strength relied exclusively on post-synaptic neurons, thus creating repeated columns in the connectivity matrix (see Okun et al., 2015, Supplementary Materials).

The prediction of this empirically confirmed cortical correlation structure supposes a further sign of the predictive power of the theory presented so far.

## 3. Discussion

Up to now, there have been presented theoretical results, based on rigorous mathematical tools, which have achieved the following goals: to give sufficient conditions for the formation of invariant, stable and low-dimensional manifolds in firing rate models, consisting on restricting the connectivity arrays to low-rank structured matrices; to prove these models can both implement, with arbitrary precision, input-driven control dynamical systems; to go through some of the theoretical consequences of the previous results, presenting propositions concerning the production of muscular activity patterns, the infinite time approximation near attracting states and the virtual power of universal computation, all of this within the predicted neural manifolds; and finally, to show how low-rank connectivity could provide circuits of a useful regularization strategy, and to present how this hypothesis truly predicts empirically validated results concerning cortical correlations.

When it comes to the network setup, a strong link between universal approximation capabilities and neural manifolds have been revealed, showing that a condition guaranteeing the universal implementation of flows is also a signature of low-dimensional dynamics, namely, the low-rank connectivity structure.

As to the external control of the modelled behavioral and cognitive phenomena, we have used many times the hypothesis that this could often be supplied by the driven modification of the initial conditions along the manifold (Beiran et al., 2023; Remington et al., 2018). Concerning external continuous input driving, theorems 2.1 and 2.2 offer vast possibilities for the simulation of externally controlled on-line computing systems, similarly to the unstructured models of liquid state machines (Maass et al., 2002; Maass & Markram, 2004).

Some of the previous achievements have similar precedents in the literature. For instance, regarding the relation between low-rank connectivity, neural manifold dynamics and universal approximation properties, akin results were recently announced in (Beiran et al., 2021; Dubreuil et al., 2022) as well as in (Gort, 2021), annex 1. In all cases, however, these relations where only explored for the v-model, as we have no evidence concerning the existence of the same results for the u-model in the literature.

Regarding the universal approximation of dynamical systems by neural networks, similar theorems were also announced previously. For instance, in (Funahashi & Nakamura, 1993), theorem 2.2 was proven for the less general class of autonomous systems; in (Trischler & D’Eleuterio, 2016), driving trajectories where included, although these where internally generated as the model continued to be essentially autonomous; in (Chow & Li, 2000; Li et al., 2005), results allowing external driving where proven for a third class of models with less relevance in the neuroscientific research; in (Doya, 1993), a result similar to our theorem 2.1 was stated, albeit a lesser level of detail.

Comparable evidence also appeared recently in (Beiran et al., 2021), although the proof there did not integrate biases in transfer functions and had thus to rely on restricted tonic inputs. We think the incorporation of variable firing thresholds in the model is not only computationally useful, but also supported by evidence (Fontaine et al., 2014).

A question arises from the universal realization of control dynamical systems by firing rate models: in top-down studies where RNN were optimized to perform animal behavioral tasks (Chaisangmongkon et al., 2017; Hennequin et al., 2014; Mante et al., 2013; Sussillo et al., 2015), was the similarity between the simulations and the data due to a robust physiologic correspondence between computational models and the targeted neural systems, or was it rather that both biological and artificial systems had adopted similar computational strategies in order to perform the same task? After all, universal simulation of dynamical systems applies under a large class of non-realistic transfer functions (in fact, it is possible for any non-polynomial mapping (Leshno & Schocken, 1993)), and thus the striking similarities between biologic and simulated neural systems could well be a matter of shared universal computational capabilities, without needing a supposedly underlying common functional structure between them.

Concerning this dilemma, bottom-up research shows that firing rate models, whenever endowed with appropriate transfer functions and biophysically based parameters, are able to replicate the frequential behavior of more realistic spiking models under different kinds of input currents (Heiberg et al., 2018; Nordlie et al., 2010; Ostojic & Brunel, 2011), supporting the hypothesis that physiological features of individual neurons uphold the emergent computational phenomena unfolding at the population level.

Considering the limitations of our approach, the low-rank hypothesis is a rather strong assumption, since the provability of a random matrix to be singular is zero. Although we have seen that this hypothesis endows neural ensembles with vast computational capabilities, it would be expected from real circuits to have full rank connection matrices, which could nevertheless lay close enough to the mentioned low-rank arrays to possess the same properties.

In this wider scenario, however, the robustness of the derived neural manifolds to small connectivity perturbations should be considered. It should also be explored the mechanisms by which neural populations could, hypothetically, implement these nearly low-rank connectomes.

It could also be noted the relative simplicity of the models studied, which did not incorporate many biophysically relevant parameters, such as synaptic time constants, in order to make the equations analytically tractable. Regarding synaptic homogeneity, however, it could be assumed that, since the approach adopted so far has effectively ignored learning mechanisms, the absence of different time constants could be a consequence of excluding synaptic potentiation from the modelled phenomena.

We think the results presented so far constitute a mathematical theory with a great explanatory power, which is consistent with many recent discoveries in systems neuroscience (Gallego et al., 2017; Sussillo, 2014; Vyas et al., 2020).

From this perspective, many internal processes controlling animal behavior, like the coordination of movements, the robust generation of central patterns or the computations mediating between stimulus and responses, are all understood as dynamical systems, externally controlled by either switching between initial conditions or by directly driving their vector fields. These continuous dynamics can be simulated by large scale neural ensembles, recurrently wired through low-rank connectivity patterns. Those mutually interacting units are capable to implement arbitrary flows through nonlinear feedback, being these dynamics parallelly and massively distributed across neural internal states, unfolding at the population level. Thus, neural correlates could not be understandable at the scale of the single neuron, whose individual activity reflects a raw mixture of the implemented dynamics, but only through the network-wide spread patterns of collective activity. These emergent motifs would, in turn, be embedded in a lower-dimensional manifold, a byproduct of the low-rank wiring, which with good reason could be considered the geometric scaffold of the implemented computations.

In particular, this paradigm possesses a great power to generate cognitive and psychologically relevant interpretations from modelled neural data. Within it, the study of the physiological basis of memory and cognition can be elegantly interpreted from a geometrical lens. For instance, concerning the study of procedural memory, the short-term storage of task-relevant information, called working memory, could be understood as temporary persistent states of the manifold dynamics, while the long-term retention of the rules governing the adaptative performance of a given task, also known as reference memory, could be inferred from the dynamics defining the input-driven transitions in neural state space (Domjan, 2010).

In the same direction, and considering proposition 2.3, the dynamics governing the patterns evolving within the neural manifold could be understood as a syntactically structured set of rules, defining computations in terms of serial manipulation of symbols, being its implementation grounded in the network wiring. This emergentist proposal is backed by the connectionist approach to cognitive science, where neurons are understood to be sub-symbolic processors from which parallelly distributed processes can be inferred. This way, neural manifolds could be considered as a point of access to these spread, somewhat blurred computations, a place from where to elucidate the mechanisms by which conceptually intelligible phenomena emerge from raw sub-symbolic neural correlates (Berkeley, 2019; Smolensky, 1988; Thomas & McClelland, 2012).

In summary, the present research has tackled the study of various neural phenomena using analytical methods, proving, on the one hand, under which conditions attracting and invariant manifolds can be found in neural systems and, on the other hand, how these conditions are linked to the emergence of universal computational capabilities. In doing so, it has been possible to generalize previous results to a wider range of firing rate models, as well as to extend such developments to more general frameworks of driven non-autonomous dynamics. Consequently, statements concerning the topological equivalence of various firing rate models have been achieved and, in addition, falsifiable predictions have been generated and subsequently contrasted, satisfactorily, with existing empirical data. All the above represents a contribution to strengthen our analytic understanding of the equations ruling firing rate networks, as well as to build a theoretical framework from which geometric and computational aspects of neural dynamics are inseparably understood.

## 5. Methods

Throughout this section, we will often consider equations (1.1) and (1.2) in vector notation, defining the vector transfer function *F*: ℝ^*N*^ → ℝ^*N*^ as *F*(*x*) = (*F*_1_(*x*_1_), …, *F*_*N*_(*x*_*N*_))^*T*^. Then, if *W, ω* are the connectivity matrices, the equations for the u-model and the v-model are, respectively,

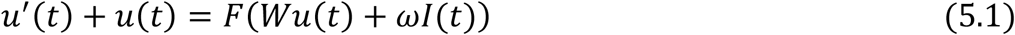

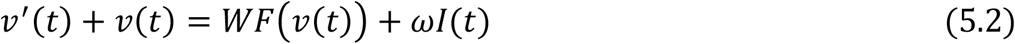

In the following sections, all the results presented in the text are going to be proved using diverse mathematical techniques from dynamical systems theory, topology and real analysis. Before every proof the most crucial results concerning these mathematical methods are going to be presented. However, some basic knowledge about the previous subjects will be assumed, and so the reader will be redirected to introductory textbooks on these topics whenever it is necessary.

### 5.1. Proof of theorem 1.1, theorem 1.2 and proposition 1.1

Throughout this paper, the notions of invariant and attracting sets have been used many times. We now formalize these concepts:

#### Definition 5.1.1

Let *x*(*t*) be some solution of a continuous, finite dimensional dynamical system for a given initial condition, *x*_0_. We say a set *S* is invariant if *x*(*t*) ∈ *S* ∀*x*_0_ ∈ *S, t* ∈ ℝ; we say a set *S* is positively invariant if *x*(*t*) ∈ *S* ∀*x*_0_ ∈ *S, t* ∈ ℝ^+^.

#### Definition 5.1.2

Let *p* ∈ ℝ^*n*^, *S* ⊂ ℝ^*n*^; Let 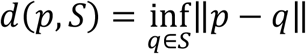; Let *x*(*t*) be some solution of a continuous, finite dimensional dynamical system for a given initial condition, *x*_0_. We say *S* is globally attracting if 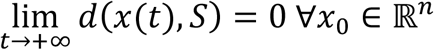.

The notion of manifold has also appeared ubiquitously in the text. We present here its rigorous definition:

#### Definition 5.1.3

A *n*-dimensional manifold is any Hausdorff, second-countable topological space, *X*, for which for every point *x* ∈ *X* there exists a neighbourhood homeomorphic to some open set of the Euclidian space, ℝ^*n*^.

Here it is not intended to make a comprehensive introduction to manifold topology, so we refer the reader to any elementary textbook on topology covering manifolds, like (Munkres, 2002), for a thorough exposition of these and other topics. Nonetheless, we present here the notion of homeomorphism, as it is going to be fundamental.

#### Definition 5.1.4

Let *X, Y* be topological spaces. A map *h*: *X* → *Y* is a *homeomorphism* whenever it is continuous, invertible and have a continuous inverse.

Homeomorphic sets can be considered as equivalent in a topological sense, since each neighbourhood in one space will be mapped to a unique neighborhood of the other, and vice versa. It is thus that homeomorphisms are regarded as topological isomorphisms.

To prove the mentioned results, we present the following lemmas:

#### Lemma 5.1.1

Suppose *W* ∈ ℝ^*N*×*N*^. Then, *rank*(*W*) = *n* if and only if there exist some rank *n* matrices *B* ∈ ℝ^*N*×*n*^, *A* ∈ ℝ^*n*×*N*^ such that *W* = *BA*

**Proof:** If *W* = *BA*, the columns of *W* are all linear combinations of the *n* linearly independent columns of *B*. Since *rank*(*A*) = *n*, the columns of *W* span the same column space of *B*, proving the forward result; If *rank*(*W*) = *n*, suppose {*b*_1,_ …, *b*_*n*_} is a basis for the image of *W*. Then, there exist unique scalars *a*_*ij*_ such that 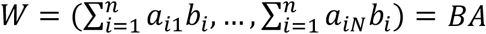, where the columns of *B* are given by *b*_*i*_ and the elements of *A* by *a*_*ij*_. By the fundamental theorem of linear algebra, *rank*(*A*) = *rank*(*W*) = *n*. This concludes the proof ▪

#### Lemma 5.1.2

Suppose {*f*_*n*_} is a sequence of continuous functions of the form *f*_*n*_: ℝ^*n*^ → ℝ, and suppose it is Cauchy under the uniform norm, meaning that

∀*ε* > 0, ∃*n*_0_ ∈ ℕ such that |*f*_*n*_(*x*) − *f*_*m*_(*x*)| < *ε* ∀*n, m* ≥ *n*_0_, *x* ∈ ℝ^*n*^ Then, there exists a continuous function *f* fulfilling that {*f*_*n*_} → *f* uniformly, this is,

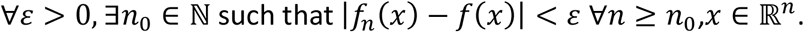

For a proof of this important result, see any textbook on real analysis covering successions and series of functions, such as (Ortega Aramburu, 2002; Rudin, 1976). We now turn to prove the theorems of section 1.1.

#### Proof of theorem 1.1

If *rank*(*W*) = *N*, it is verified trivially that ℳ = ℝ^*N*^. If, for the contrary, *rank*(*W*) = *n* < *N*, we know from lemma 5.1 that *W* = *BA*, where both the columns of B and the rows of A are linearly independent vectors. We are going to perform the change of variables 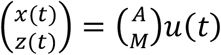, where the matrix *M* has the *N* − *n* basis vectors of the kernel of *W* as its rows.

In this new basis, the system can be expressed as:

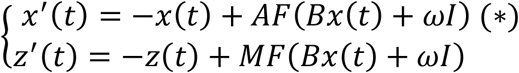

where *I* is, by hypothesis, a constant input. Let’s see, for later, that if 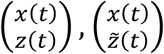 are both solutions of the previous system of ordinary differential equations (ODEs), then

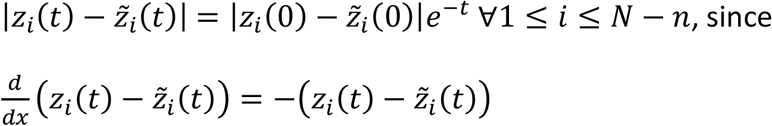

We will now focus on the study of the system of ODEs given by (*) plus the i-th component of *z*(*t*), which is passively driven by *x*(*t*). We will label this *n* + 1 dimensional autonomous system by (**), and their solutions are going to be represented as 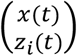.

Now, we proceed to define some sets: let 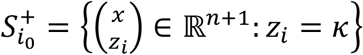, where *κ* ≔ ∑_*i,j*_|*m*_*ij*_|, being *m*_*ij*_ the components of *M*; if *M*_*i*_ is the i-th row of *M*, then let

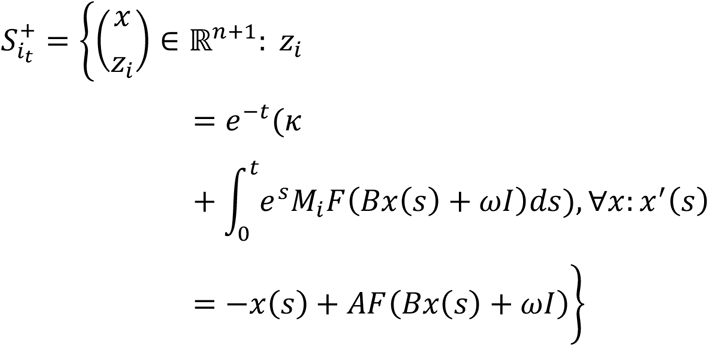

so that the previous set is the image of 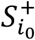 under the flow of (**) after a time *t*; we can define, analogously, 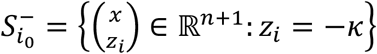,and 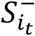, in the same way, making again the change *κ* ↦ −*κ* ; finally, we define the set 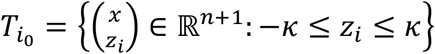, so that 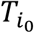 is the closed set which includes the origin and whose frontier is 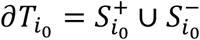.

It can be checked that 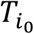 is positively invariant. Indeed, *z*_*i*_(−*z*_*i*_ + *M*_*i*_*F*(*Bx* + *ωI*)) < 0 for every 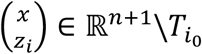, since, by definition, *F* is bounded by 1. Thus, no trajectory can escape 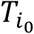, as, by the mean value theorem, the contrary would imply the existence of a time *c*: *z*_*i*_(*c*) *z*_*i*_′(*c*) > 0. A similar argument can be used to show that 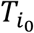 is globally attracting.

We now prove that, for all 0 ≤ *t*, there exists a continuous function 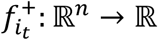 fulfilling that

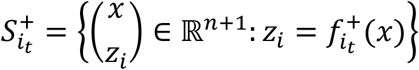

Indeed, given *x*_0_ ∈ ℝ^*n*^, let *x*(*t*) be the solution of (*) with *x*(0) = *x*_0_, which exists and is unique for all ℝ, since the system (*) is globally Lipschitz. Then: 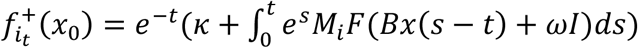. The fact that 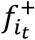 is continuous comes from the continuity of the flow. Analogously, we can also define continuous functions 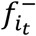 such that

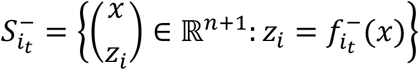

Let’s now see that 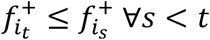. If the contrary held, ∃*x*_0_ ∈ ℝ^*n*^, 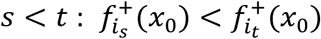. In this case, let *x*(*t*) be the solution of (*) with *x*(0) = *x*_0_, and *z*_*i*_(*t*), 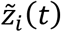 be such that 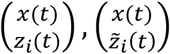 are both solutions of the *n* + 1 dimensional system, (**). Suppose further that 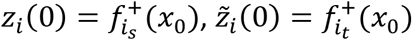. Then, by the definition of 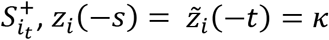. But since we showed 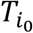 is positively invariant, 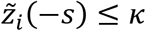. Thus, using Boltzano’s theorem, ∃*c* ∈ [−*s*, 0) such that 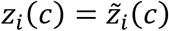. This, however, contradicts Picard’s existence and uniqueness theorem, thus proving our claim that the functions 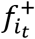 are monotonically decreasing. Using the same procedure, it can be shown that 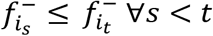, being this complementary set of functions monotonically increasing.

We now define a sequence of functions, 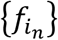, given by 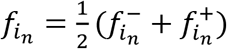. We claim this sequence is Cauchy under the uniform norm. To see this, let *ε* > 0 be given; choose a natural number 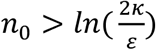 so that 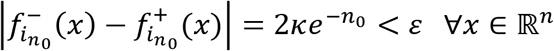, using what we proved earlier. Because of monotony, 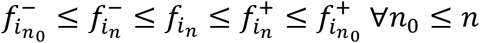, and therefore

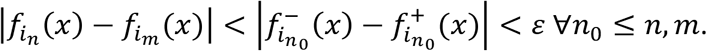

Since 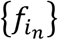 is a Cauchy sequence, lemma 5.2 stablishes the existence of a continuous function, *f*_*i*_, such that 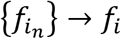 uniformly.

Define 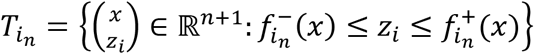, which forms a sequence of nested closed sets, 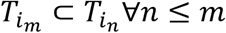. Notice that every 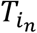 is positively invariant and globally attracting given the flow of (**), since each of them is an image of 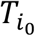 under the flow map. Moreover, 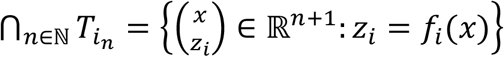, since 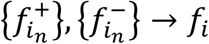 uniformly. With all this, the set 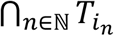 is found to be positively invariant and globally attractive in the *n* + 1 dimensional dynamical system given by (**). The fact that this set is also invariant for all backward time comes from the fact that 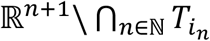 is disconnected, being it the union of two connected invariant sets.

To finish the proof, we repeat the previous reasoning for every 1 ≤ *i* ≤ *N* − *n*, finding that the set 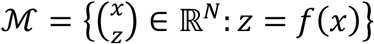 is invariant and globally attracting in the original *N* dimensional system, where *f*(*x*) = (*x*_1_, …, *x*_*n*_, *f*_1_(*x*), …, *f*_*N*−*n*_(*x*)). Since *f*: ℝ^*n*^ → ℳ is a homeomorphism, ℳ is also a manifold. This concludes the proof ▪

#### Proof of theorem 1.2

Again, if *rank*(*W*) = *N* it is verified trivially that our Euclidean manifold is given by ℝ^*N*^. If *rank*(*W*) = *n* < *N*, we decompose *W* = *BA*. It is going to be proven that ℳ = {*v* ∈ ℝ^*N*^: *v* = *Bx* + *ωI, x* ∈ ℝ^*n*^} is globally attractive and invariant. Let *M* be the matrix whose rows span the normal space of ℳ, so that *MB* gives the null (*N* − *n*) × *n* matrix. Let *v*(*t*) be any solution of the v-model. Then: 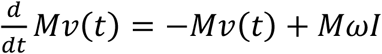. Solving this linear differential equation, we find that *Mv*(*t*) = (*Mv*(0) − *MωI*)*e*^−*t*^ + *MωI*. Therefore, for any *t* ∈ ℝ: *v*(*t*) ∈ ℳ ⟺ *M*(*v*(*t*) − *ωI*) = 0 ⟺ *M*(*v*(0) − *ωI*) = 0 ⟺ *v*(0) ∈ ℳ With this we proved ℳ is invariant. To show it is also globally attractive it is enough to see that, for any initial condition, 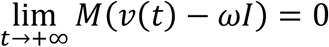▪

For proposition 1.1, we define the notion of topological conjugacy:

#### Definition 5.1.5

Suppose we have two *C*^1^ finite-dimensional dynamical systems, and suppose there exists a homeomorphism, *h*, such that, for any solution of the first system, *x*(*t*), we have that *h*(*x*(*t*)) is also a solution of the second one. Then, we say that these dynamical systems are *topologically conjugate*. The homeomorphism, *h*, is called the conjugacy.

A more general definition, which the previous one can be shown to fulfil, would be to say that two dynamical systems are conjugate whenever there exists a conjugacy which commutes with the flow map. However, the previous definition is enough for the scope of this work. With this, we are ready to prove the last result of section 1.1, which will rely heavily on the previous proofs.

#### Proof of proposition 1.1

Suppose *u*(*t*), *v*(*t*) are solutions of the u-model and the v-model, respectively. In the case where *rank*(*W*) = *N*, choose *h*: ℝ^*N*^ → ℝ^*N*^: *h*(*x*) = *Wx* + *ωI* to be the conjugacy, and verify that, for every solution *u*(*t*) of the u-model, we have that

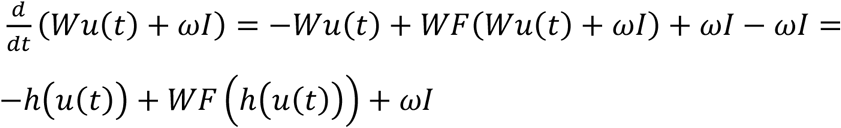

Thus, *v*(*t*) = *h*(*u*(*t*) is a solution of the v-model, and therefore *h* is a conjugacy.

In the case of *rank*(*W*) = *n* < *N*, decompose *W* = *BA*. Without loss of generality, we can assume the columns of *B* form an orthonormal basis of the image of *W*. Let’s call 𝒩 to the *n*-dimensional Euclidean manifold obtained in theorem 1.2, and *x* to the *n*-dimensional vector whose elements define the coordinates of 𝒩 in the base given by *B*. Then, *v*(*t*) ∈ 𝒩 ⇔ *v*(*t*) = *Bx*(*t*) + *ωI*. Since *x*(*t*) = *B*^*T*^(*v*(*t*) − *ωI*), we have that any *x*(*t*) is the solution of

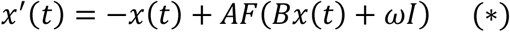

This defines the neural manifold dynamics in 𝒩.

Now consider any solution, *u*(*t*), on the u-model’s neural manifold, ℳ, and decompose *W* = *BA* in the same way. Suppose that *x*(*t*) is some solution of (*), and perform a change of basis such that 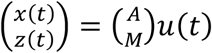, where the matrix *M* has a basis of *W*^′^s kernel as it’s rows, as done in the proof of theorem 1.1. There, it was proved that the solutions of the u-model in their neural manifold, ℳ, are given by

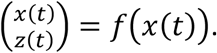

To finish the proof, define the homeomorphism:

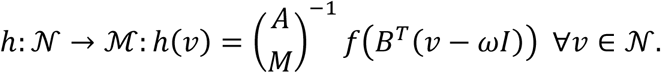

It is now straightforward to see that, for every solution *v*(*t*) ∈ 𝒩 of the v-model, we have a solution *h*(*v*(*t*)) ∈ ℳ of the u-model, as we wanted.

▪

### 5.2. Proof of theorems 2.1 and 2.2

In this subsection, theorems expressing the universal approximation capabilities of firing rate models are going to be proved. For this reason, we present the following preliminary work:

#### Theorem 5.2.1

**(approximation by sigmoidal superposition):** Let *K* be a compact set of ℝ^*n*^, and *f*: *K* → ℝ^*m*^ a continuous function. Then, given an arbitrarily small *ε* > 0, there are *N* ∈ ℕ and matrices *A* (*m*× *N*), *B* (*N* × *n*), *θ* (*N* × 1) such that:

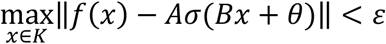

where *σ*: ℝ → (0, 1) is a *sigmoid map*, this is, a monotonically increasing continuous function such that 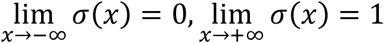. The theorem also holds for more general functions, as long as they are non-polynomic (Leshno & Schocken, 1993).

For proofs of this important result in neural network theory, see (Cybenko, 1989; K. I. Funahashi, 1989; Tinchcombe et al., 1989).

In the following proofs, we will need a stronger definition of continuity:

#### Definition 5.2.1

Let *U* ⊂ ℝ^*m*^. We say a function *f*: *U* → ℝ^*m*^ is Lipschitz continuous if there exists a constant *C* > 0 such that ‖*f*(*x*) − *f*(*y*)‖ ≤ *C*‖*x* − *y*‖∀*x, y* ∈ *U*; we say a function defined as before is locally Lipschitz if for every *p* ∈ *U* there exists a neighborhood of *p, U*_*p*_, where *f*|*U*_*p*_ is Lipschitz continuous.

#### Lemma 5.2.1

If a function is differentiable, then it’s locally Lipschitz; if a function is locally Lipschitz, it is Lipschitz in every compact set *K* ⊂ *U*. For a definition of compactness, see the following subsection.

#### Lemma 5.2.2

Let *U* ⊂ ℝ^*n*^ × ℝ be an open set containing (*x*_0_, 0) and suppose that the maps *F*, 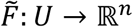 are locally Lipschitz. Suppose also that 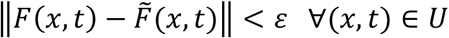 for some *ε* > 0. Let *C* be a Lipschitz constant in *x* for *F*. If *x*(*t*), *y*(*t*): *I* ⊂ ℝ → ℝ^*n*^ are solutions, respectively, of the systems of equations given by *x*^′^(*t*) = *F*(*x*(*t*), *t*), 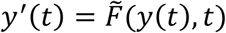, on some interval *I* and these solutions fulfil that *x*(0) = *y*(0) = *x*_0_, then:

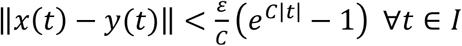

For the proofs of the previous lemmas, we redirect the reader to (Hirsch et al., 2013b).

#### Lemma 5.2.3

let *v*(*t*) ∈ ℝ^*n*^ solution of the system

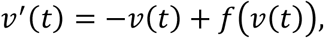

where *f*: ℝ^*n*^ → ℝ^*n*^ is Lipschitz, and *u*(*t*) ∈ ℝ^*n*^ be solution of *u*^′^(*t*) = −*u*(*t*) + *f*(*v*(*t*)). Then, *u*(0) = *v*(0) ⇒ *u*(*t*) = *v*(*t*) ∀*t* ∈ ℝ

**Proof:** Knowing that solutions are unique and defined for all time, it’s only left to show that, for all the components of the solution vectors:

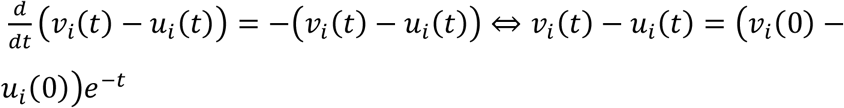

#### Proof of theorem 2.1

Let *ε, T* > 0 be given. Let *x*(*t*) be the solution of the initial value problem of the system to be approximated for some *x*(0) = *x*_0_ ∈ Ω, and suppose [0, *T*] is included in its maximal interval. Let 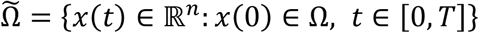 be the image of Ω under the flow map for the time interval [0, *T*].

Start supposing that *W* = *BA*, being *B, A*, matrices as the ones described in lemma 5.1.1, whose precise value is going to be provided later. Define *y*(*t*) ≔ *Au*(*t*), so that *y*^′^(*t*) = −*y*(*t*) + *AF*(*By*(*t*) + *ωI*(*t*)). Let *C*_*G*_ be a Lipschitz constant for *G* in Ω, whose existence is guaranteed since *G* is assumed to be differentiable, and thus locally Lipschitz (lemma 5.2.1). Then, theorem 5.2.1 guarantees the existence of *N* ∈ ℕ and matrices *ω*(*n* × *N*), *A* (*n* × *N*), *B* (*N* × *n*), such that:

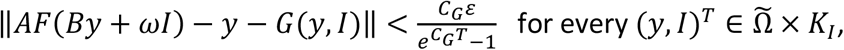

whenever *F* is endowed with appropriate thresholds. Here,

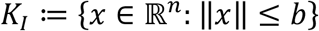

where *b* is taken to be big enough so that *I*(*t*) ∈ *K*_*I*_ ∀*t* ∈ ℝ^+^ for all input trajectories drawn from the uniformly bounded set of functions considered in the statement of the theorem.

Choose *u*(0) such that *Au*(0) = *x*_0._This system is solvable since, as stated earlier, we can assume that *rank*(*A*) = *n*, as these matrices form an open and dense set in ℝ^*n*×*N*^. Then, by lemma 5.2.2:

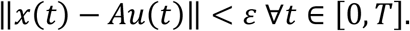

Letting *R* = *A*, we found what we wanted to prove. Since *W* = *BA*, lemma 5.1.1 guarantees that *rank*(*W*) = *n*. ▪

#### Proof of theorem 2.2

Let *ε, T* > 0 be given. Let *x*(*t*) be the solution of the initial value problem of the system to be approximated for some *x*(0) = *x*_0_ ∈ Ω, and suppose [0, *T*] is included in its maximal interval. Let 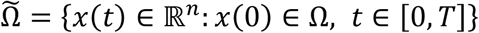, *t* ∈ [0, *T*]} be the image of Ω under the flow map for the time interval [0, *T*].

Define *y*(*t*) ∈ ℝ^*n*^ as the solution of the initial value problem given by

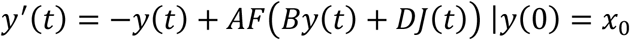

Where *A* is some *n* × (*N* − *n*)matrix, *B* is (*N* − *n*) × *n* and *D* is of order (*N* − *n*) × *m*. Let *C*_*G*_ be a Lipschitz constant for *G* in Ω, whose existence is guaranteed since *G* is assumed to be differentiable, and thus locally Lipschitz. It is clear that 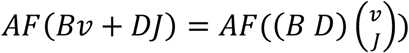. With this, we can make use of theorem 5.2.1, which assures that there exist matrices *A*, (*B D*) as the ones we just defined such that, for *N* sufficiently big, ratify that:

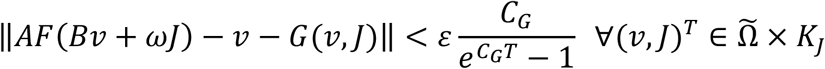

whenever *F* is endowed with appropriate thresholds. Here,

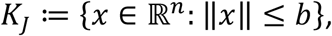

where *b* is taken to be big enough so that *J*(*t*) ∈ *K*_*I*_ ∀*t* ∈ ℝ^+^ for all the possible driving trajectories, *J*, drawn from the uniformly bounded set of functions considered in the statement of the theorem.

With all the previous, lemma 5.2.2 establishes that

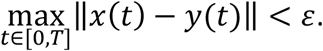

Let’s define the connectivity matrices of the net, *W* (*N* × *N*), *ω*(*N* × *m*), as 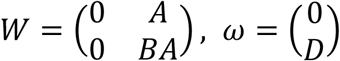; we will split the solutions of the v-model, making *v*(*t*) = (*v*^1^(*t*), *v*^2^(*t*))^*T*^, where *v*^1^(*t*) ∈ ℝ^*n*^, *v*^2^(*t*) ∈ ℝ^*N*−*n*^.

Now, we check that *v*^2^(*t*) = *By*(*t*) + *DJ*(*t*) is the value of the last components of the solution vector, *v*(*t*), whenever *v*^2^(0) = *Bx*_0_ + *DJ*(0), since:

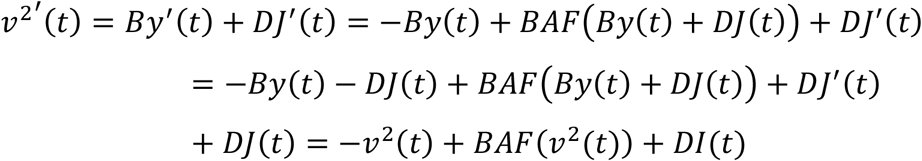

Using that, by definition, *I*(*t*) = *J*′(*t*) + *J*(*t*). Now, since the equations of the first components of *v*(*t*) are given by *v*^1^′(*t*) = −*v*^1^(*t*) + *AF*(*v*^2^(*t*)), choosing *v*^1^(0) = *x*_0_ we have, by lemma 5.2.3, that *v*^1^(*t*) = *y*(*t*). We define *R* as the matrix fulfilling that *Rv*(*t*) = *v*^1^(*t*). Therefore:

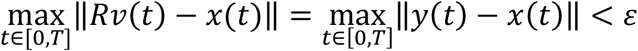

To finish the proof, we see that 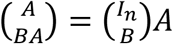, where *I*_*n*_ is the n-th order identity matrix, and thus, by lemma 5.1.1, *rank*(*W*) = *n*. With this, everything that was claimed in theorem 2.2 has been proved. ▪

### 5.3. Proof of propositions 2.1 to 2.3

We start by presenting a handful of preliminary concepts that will be needed for the proof of proposition 2.1. The notion of compacity is of special interest in the fields of topology and analysis, and we have used it implicitly in the previous results. Because in the next proof we will need a more concise grasp of this concept, we present it here its formal definition.

#### Definition 5.3.1

Let *X* be a topological space and let *K* ⊆ *X*. Suppose {*U*_*i*_}_*i*∈*I*_is a family of open sets indexed by some set *I*. We say it is an open cover of *K* if *K* ⊂ ⋃_*i*∈*I*_ *U*_*i*_; We say 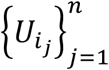 is a finite subcover of *K* if 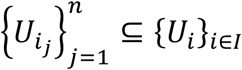 and 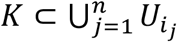; finally, we say that *K* is compact if for every open cover there exists a finite subcover for *K*.

In Euclidean spaces, the Heine-Borel theorem assures that a set is compact if and only if it is closed and bounded. Although the previous are necessary conditions in general topological spaces, they are not sufficient in more general topological spaces. This is the case in function spaces equipped with the uniform topology, as the one we deal in proposition 2.1, where distances are induced by the norm 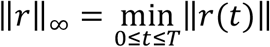. The existence of compact sets in these Banach spaces is guaranteed by the Arzelà-Ascoli theorem (Munkres, 2002). We now present an extension theorem:

#### Theorem 5.3.1

**(Kirszbraun):** Suppose *K* ⊂ ℝ^*n*^ is a compact set. Suppose also that *f*: *K* → ℝ^*m*^ is Lipschitz continuous, with a given constant *C* > 0. Then, there exists a function 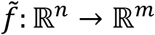 with the same Lipschitz constant, *C*, such that 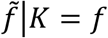

For a recent constructive proof of this theorem, see (Azagra et al., 2021). Finally, we have the following lemma:

#### Lemma 5.3.1

Under the uniform norm, *C*^∞^ functions are dense in the Banach spaces of continuous functions with compact domain.

The proof is a straightforward application of the Stone-Weierstrass theorem. See (Ortega Aramburu, 2002; Rudin, 1976) for a statement of this crucial result in mathematical analysis.

#### Proof of proposition 2.1

Considering the uniform norm, suppose 𝒜 is a compact set of continuous functions of the form *r*: [0, *T*] → ℝ^*n*^, *T* > 0, as the statement of the proposition suggests. Let *ε* > 0 be given. Let 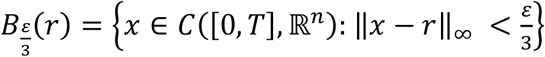 be the ball of radius 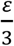 centred at *r*. Since 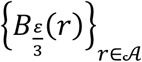 is an open cover of the compact set 𝒜, there exist a finite set of trajectories, {*r*_1_, …, *r*_*l*_} ⊆ 𝒜 for which the inclusion 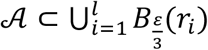 holds.

We define 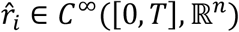 such that 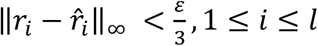, using lemma 5.3.1.

Now consider the *l* trajectories in ℝ^*n*+*l*^, 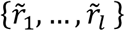, whose components are given by

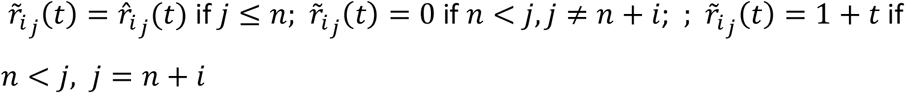

It can be checked that 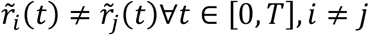, and that 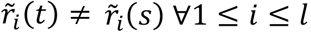, so these new trajectories will never cross each other. Now consider the compact set of ℝ^*n*+*l*^ defined as follows:

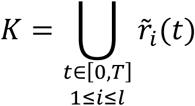

Consider also the function *g*: *K* → ℝ^*n*+*l*^ which fulfils that 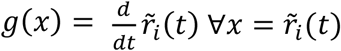. Since every 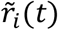 is *C*^∞^, and because these trajectories never cross, the function is well-defined. Moreover, since they all have continuous derivatives, by lemma 5.2.1 *g* is Lipschitz. Then, theorem 5.3.1 allows to extend *g* to a Lipschitz function *G*: ℝ^*n*+*l*^ → ℝ^*n*+*l*^.

Suppose we want to approximate an arbitrary trajectory *r* ∈ 𝒜. To this end, from compactness we know that ∃1 ≤ *i* ≤ *l* such that 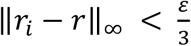. Now let *x*(*t*) be the solution of the initial value problem 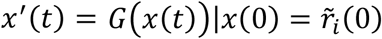. From the existence and uniqueness theorem, it follows that 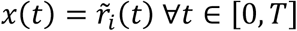

Although we presented theorems 2.1 and 2.2 in terms of dynamical systems induced by *C*^1^ functions to avoid technicalities, there is nothing restricting us to apply the same results to more general Lipschitz functions. Thus, by theorem 2.1, there exists a u-model whose output fulfils that 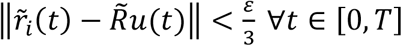. Now, let *M* ∈ ℝ^*n*×(*n*+*l*)^ be given by *M* = (*I*_*n*_ 0). Let 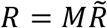. Then, using the triangle inequality:

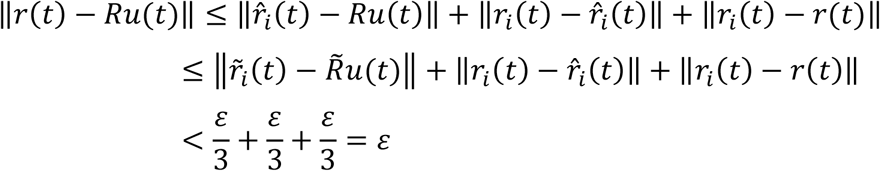

For the v-model, use instead theorem 2.2 to obtain the same result. This concludes the proof.

▪

Now we turn to the proof of proposition 2.2, which involves periodic solutions of differential equations, this is, trajectories *s* for which ∃*T* > 0: *s*(*t*) = *s*(*t* + *T*) ∀*t* ∈ ℝ. The orbits fitting in this definition consists of either fixed points or limit cycle oscillations.

The following deduction will require of new definitions and preliminary results. The first one involves the rigorous formulation of the notion of a flow, which we eluded so far in order to leave aside the more technical concepts from the main text.

#### Definition 5.3.2

Let *U* ⊆ ℝ^*n*^ be open. A map *g*: *U* × ℝ → *U* is called a *flow* if for every *s, t* ∈ ℝ and *x* ∈ *U*, it fulfils the following axioms:

- *g*_0_(*x*) = *x*
- *g*_*s*+*t*_(*x*) = *g*_*s*_(*g*_*t*_(*x*))

In the case of continuous time dynamical systems induced by Lipschitz functions, the existence and uniqueness theorem allow to express the bundle of its solutions as a flow fulfilling the equation

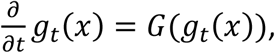

being *x* the values for the initial conditions. A remarkable property of this flow is the following:

#### Theorem 5.3.2

Let *G*: *U* ⊆ ℝ^*n*^ → ℝ^*n*^ have Lipschitz constant *C*_*G*_, and let the flow map of its induced dynamical system be given by *g*: *U* × ℝ → *U*. Then, it is fulfilled that

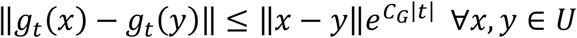

In other words, this result assures that the flow mapping is continuous. For a proof, see (Hirsch et al., 2013b).

In the case of the flow on a neural manifold, ℳ, of a low-rank system, we will design it by υ_*t*_(*f*(*x*)), where *x* ∈ ℝ^*n*^ stand for the initial position on the intrinsic coordinate system of the manifold. As it was seen in the proof of theorem 1.1 (and as it can be seen trivially in the proof of theorem 1.2), the coordinate chart of ℳ can be expressed by a continuous bijection, given by *f*: ℝ^*n*^ → ℳ, that can be chosen so that its inverse is defined by the linear surjection *R*: ℝ^*N*^ → ℝ^*n*^ restricted to ℳ, where *R* was the matrix set to define the readout of the network in theorems 2.1 and 2.2. This way, in order to study neural manifold dynamics, we assure the initial condition of the network, *f*(*x*), lays on the manifold, and thus that the correspondent solution remains there forever.

Before we start proving proposition 2.2, we present another important result concerning the topology of compact spaces:

#### Theorem 5.3.3

Let *X* be a metric space. Then, it is compact if and only if for every sequence {*x*_*n*_} ⊂ *X* there exists a subsequence 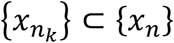 such that 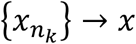, for some *x* ∈ *X*.

An interesting idea used in the proof of this theorem, which is not going to be presented here, is the notion of a Lebesgue number. Given an open cover of a compact set, there exists a number, *λ* > 0, such that the ball with radius *λ* centred in any point on the compact region is included in some of the open sets forming the cover. For a proof, see (Ortega Aramburu, 2002). Now proposition 2.2 is proved.

#### Proof of proposition 2.2

Let *ε* > 0 be given. Let *S* = ⋃_*t*∈ℝ_ *s*(*t*) be the orbit set traced by some attracting solution. We will suppose there is just one of them, as the process below can be repeated for each of the open sets composing the original set *U*, which is necessarily disconnected in the case there are many stable trajectories. Let *V* be a bounded open set such that 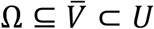, being 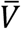 its closure. It can be constructed as the union of a set of balls with sufficiently small radius.

Let 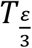 be big enough in order to satisfy that

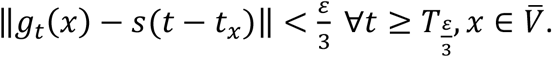

To prove the existence of such number, negate the claim to find a succession 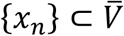 for which, for each term, 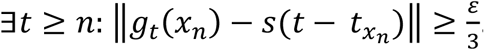. As 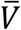 is compact, by theorem 5.3.3 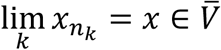. But then, by theorem 5.3.2, *g*_*t*_(*x*) should not converge to *s*(*t* − *t*_*x*_), thus obtaining a contradiction. The open set 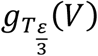 has the property that

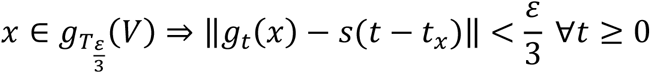

Let *λ* > 0 be such that 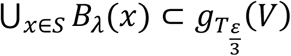. Given that 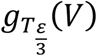 is an open cover of *S, λ* can be taken to be the Lebesgue number of that cover. It’s obvious that 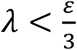.

Let 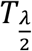 be defined analogously, so that 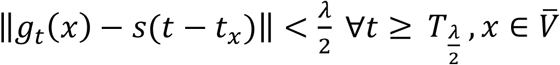. Again, the existence of such time is proved as with 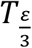. We define a function 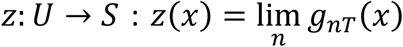, so that it represents to which phase of the attracting orbit does the solution with initial value *x* converge. To conclude this list of definitions, let *W*_*x*_ = {*y* ∈ *U*: *z*(*y*) = *z*(*x*)}, so that it is the set of initial values which converge to the same orbit with the same phase.

First, we prove *z*(*x*) is continuous. To this end, let *η* > 0 be given, and let *n* ∈ ℕ such that 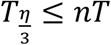, where 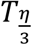 is defined as in the previous cases. Then, by theorem 5.3.2:

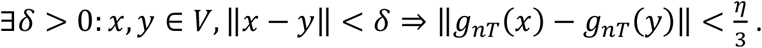

But then, by the definition of 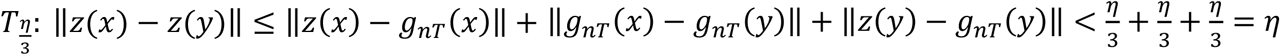

Now, by theorem 2.1 (resp. theorem 2.2) it can be built a u-model (resp. a v-model) whose flow fulfils that 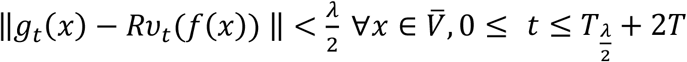. From the triangle inequality, it follows that ‖*s*(*t* − *t*_*x*_) − *R*υ_*t*_(*f*(*x*)) ‖ ≤ ‖*s*(*t* − *t*_*x*_) − *g*_*t*_(*x*)‖ + ‖*g*_*t*_(*x*) − *R*υ_*t*_(*f*(*x*)) ‖ < *λ* for every 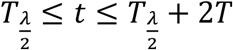.

In the limit cycle case, without loss of generality, we can assume that *λ* is small enough so that *S* ⊈ *z*(*B*_*λ*_(*s*(*t*)))∀*t* ∈ ℝ. Since 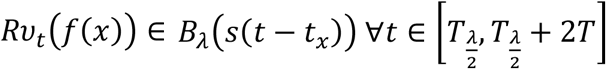, parametrizing *S* and using the intermediate value theorem, it can be shown that there exists some value of time 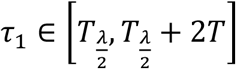 where 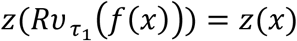. The same result is evident in the fixed-point scenario. Thus, in both situations one has that 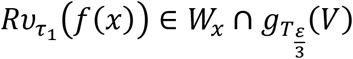.

Let *n* ∈ ℕ and suppose ∃*τ*_*n*_ ∈ ℝ^+^ such that 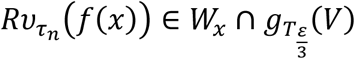.

Then:

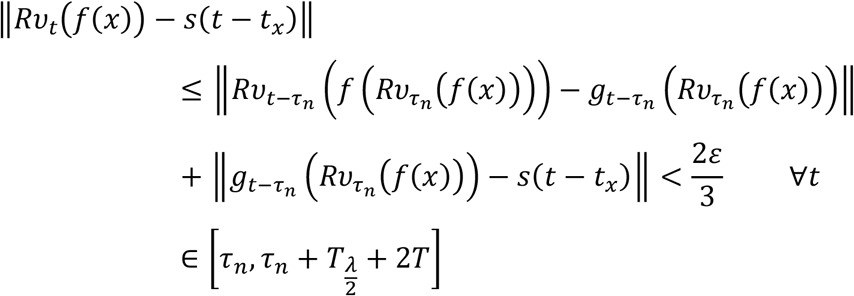

In addition, repeating the argument used for *τ*_1_, there exists a 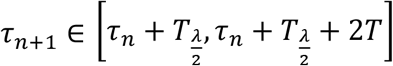 such that 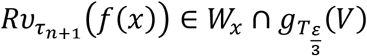. Thus, by induction, we have, for all 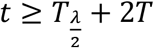:

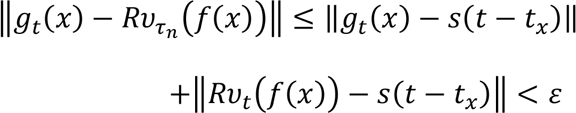

To finish this subsection, the proof of proposition 2.3 is going to be presented, relating universal approximation capabilities of neural networks to the simulation of symbolic computations whenever finite (yet arbitrary) memory is required. From now on, we will follow the Church-Turing thesis so that for any decidable function, *f*, we will assume that an effective procedure capable of computing its outputs can be implemented by a Turing machine. In order to link these theoretical devices with dynamical system theory, we present the following theorem, which claims that discrete time dynamical systems possess the power of universal computation.

#### Theorem 5.3.4

Every Turing machine is equivalent to an iterated map of the form *F*: ℕ → ℕ.

**Proof:** Describe the configuration of a given Turing machine as follows: suppose that {*i*_*n*_} is the sequence describing the tape, being each term the number codifying each symbol of the tape once a particular order has been assigned to the alphabet set, where *i* = 0 stands for the blank symbol, the only appearing infinitely many times; {*p*_*n*_} = {2, 3, 5, 7, 11 …} is the sequence of prime numbers; *s* ∈ ℕ the state, being *s* = 1 the codification of the initial state; *N* ∈ ℕ the position on the tape. Then, the natural number

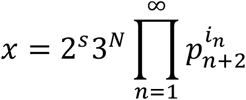

codifies uniquely the configuration of the Turing machine. Thus, it can be constructed a function *F*: ℕ → ℕ implementing the transition function with which the given Turing machine operates.

▪

However, concerning neural computations, since the models studied in this paper take continuous values of both time and phase variables, it can be questioned how these kinds of systems could effectively implement an essentially discrete model of computation. It is thus that we should define in which way it could be understood that a continuous system simulates a discrete one. For that purpose, we present some theory developed by Branicky in (Branicky, 1995).

#### Definition 5.3.3

**(S-simulation):** Let *X, Y* be topological spaces. Let *g*: *X* × ℝ → *X* be the flow of a continuous dynamical system, and *F*: *Y* → *Y* an iterated map describing a discrete one. We say the former system S-simulates the later whenever there exists a continuous surjective function *ψ*: *D* ⊆ *X* → *Y* and some *T*_0_ ∈ ℝ^+^ such that

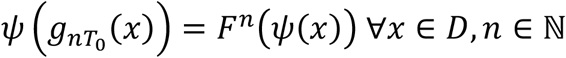

where *F*^*n*^ denotes the map *F n* times composed with itself.

This definition is similar to definition 5.1.4, which presented the notion of topological conjugacy. In this case, however, the function *ψ* needs not to be injective, and thus topological conjugacy is a stronger hypothesis compared to S-simulation.

With this, we could set up a neural simulation of some Turing machine as follows: choose some intrinsic dimensionality for the neural manifold (it is going to be proved that 3 dimensions are sufficient) and place a collection of open sets in this immersed space, each one of which will stand for a particular configuration of the Turing machine’s tape, position and current state. Then, use theorems 2.1 and 2.2 to define a dynamical system capable to implement the transition function of the algorithm. This could be performed by joining the mentioned open regions with concrete phase trajectories, indicating the system which steps should it follow to simulate the given computation. In order to formalize the previous intuition, we present the following theorem.

#### Theorem 5.3.5

Let *F*: *Y* ⊂ ℤ^*n*^ → ℤ^*n*^ define a discrete dynamical system and suppose *Y* is a compact subset. Then, it can be S-simulated by a continuous-time dynamical system in ℝ^2*n*+1^ whose dynamics are induced by a Lipschitz function.

For a proof see (Branicky, 1995), theorem 5.7.

It is interesting to see that this dynamical system is robust, being able to carry out the simulation also in the cases where the flow is slightly perturbed or when some small enough amount of noise is added to the model.

Indeed, in the original proof (Branicky, 1995) it was constructed a continuous nearest integer function like

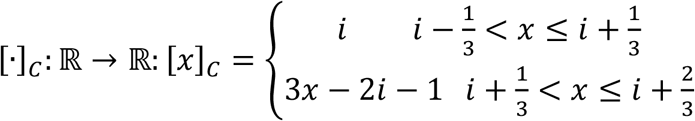

for all *i* ∈ ℤ, subsequently defining the map inducing the S-simulation, *ψ*: ℝ^2*n*+1^ → ℝ^*n*^, like *ψ*(*x*) = ([*x*_1_]_*C*_, …, [*x*_*n*_]_*C*_) ^*T*^∀ *x* ∈ ℝ^2*n*+1^.

Now suppose that *g*_*t*_(*x*) is the flow simulating the given iterated map. To say that this system S-simulates a given discrete time dynamical system is equivalent to say that for every initial condition *x* ∈ *ψ*^−1^(*F*(*Y*)) it is fulfilled that 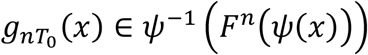 for all *n* ∈ ℕ. It can be seen from the definition of *ψ* that *ψ*^−1^(*Y*) has nonempty interior, since for every *y* ∈ *Y*, if *x* = (*y, y*, 0)^*T*^ then 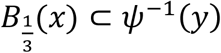. Therefore, given any *M* ∈ ℕ, for every 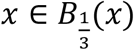 one can find some *r* > 0 such that 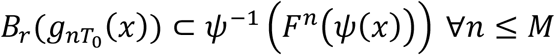. This is saying that if some small enough perturbation is added to some trajectory starting in an open set, it will still be able to efficiently carry on the computing process during a sufficiently long period of time, thus ensuring that our setup is robust. In the proof of proposition 2.3, which is presented below, it is shown how robustness is preserved by neural manifold implementations.

#### Proof of proposition 2.3

By the Church-Turing thesis, take the effective procedure which sustain the computable function *f* to be performed by some Turing Machine. Let *S* be a set of natural numbers, each of which codifying a given tape uniquely as it was done in the proof of theorem 5.3.4. Since we restrict our computations to require only a finite amount of memory, there exists *l* ∈ ℕ such that the length of the nontrivial segment of the tape is less than *l* for every tape codified in *S*. Let *M* ∈ ℕ be the maximum number of epochs necessary to terminate the program in any input of the given set, *S*.

By theorem 5.3.4, there is a map *F*: *Y* ⊂ ℕ → ℕ implementing the computation, where we define the set 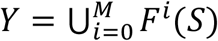. It is compact since it is a finite set of natural numbers, each of which represents some configuration of the Turing machine. It is invariant since after a time *M* every possible computation has been completed, reaching thus a stable state of the mapping.

By theorem 5.3.5, there exists a Lipschitz flow *g*: ℝ^3^ × ℝ → ℝ^3^ S-simulating the dynamical system produced by *F*, where the function *ψ* defining the simulation can be the one we already presented.

Then, choose some 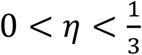 and let Ω = {*x* ∈ ℝ^3^: ‖*x* − *y*‖ ≥ *η* ∀*y* ∈ ℝ^3^\*ψ*^−1^(*S*)}. It can be seen that Ω ⊂ *ψ*^−1^(*S*)°, where *ψ*^−1^(*S*)° denotes the interior of *ψ*^−1^(*S*), and that Ω is compact.

Let 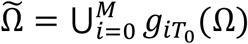, which is still compact. Since *g* S-simulates the discrete dynamics, it follows that 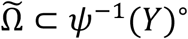, where ° stands again for the interior. This is because the Lipschitz condition makes 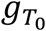 a homeomorphism, and thus it maps interiors to interiors.

Since *ψ*^−1^(*Y*)° is an open cover of 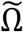, let *λ* > 0 be a Lebesgue number. Using theorem 2.1 in the u-model case or theorem 2.2 for the v-model, we have can find a recurrent network such that

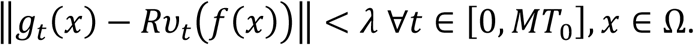

With this, it is seen from the definition of 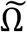 that 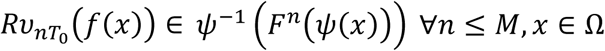, *x* ∈ Ω, which implies that

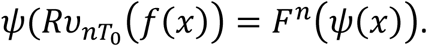

It only remains to be shown that the neural manifold of this system is 3-dimensional. Indeed, theorems 5.3.4 and 5.3.5 together predict that *g* can be a 3-d flow. Thus, using theorems 2.1 and 2.2 *rank*(*W*) = 3 in both models. Therefore, theorems 1.1 and 1.2 confirm that the emerging manifolds will be 3-dimensional. This concludes the proof. ▪

Using wat it is presented in the previous proof, since Ω has nonempty interior, if *x* ∈ Ω°, using that ‖*g*_*t*_(*x*) − *R*υ_*t*_(*f*(*x*))‖ < *λ* and the same arguments we fenced to show *g* is robust, it also follows that the computational process held by the neural manifold trajectory υ_*t*_(*f*(*x*)) will tolerate small perturbations without affecting its performance, both if we slightly modify the trajectory or if we do so for its initial condition by a sufficiently small amount.

### 5.4. Descriptive statistics of low-rank models

In this subsection a precise formulation of the statement presented in section 2.3, where it was claimed that in low-rank wirings the correlation matrix’s number of degrees of freedom was of order N, is going to be presented. We start by making the idea of order more precise:

#### Definition 5.4.1

We say that a real valued function *f* is of order *g*, being *g* a positive valued function, whenever 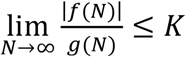, for some *K* ∈ ℝ^+^, and we write it *f*∼*O*(*g*).

The flow of the set of ODE’s defining our neural systems is going to be given by *v*: ℝ^*N*^ × ℝ^*p*^ × ℝ → ℝ^*N*^ and it is going to be written like *v*_*t*_(*x, α*), where *α* ∈ ℝ^*p*^ stand for the vector of bifurcating parameters defining the system, which include the weights, the firing thresholds or other scalars that could be added to improve the model’s fitting to data, such as membrane time constants or parameters describing different input-spiking rate relations.

Let’s now describe the statistical measures which are going to be required: *E*[*v*_*t*_(*x, α*)_*i*_] will stand for the mean of the i-th component of the trajectory starting in *x* with bifurcation vector *α* during some arbitrary lapse of time [0, *T*], *T* > 0, and will be given by

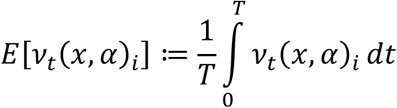

Taking the previous definition, *C*_*ij*_(*x, α*) ≔ *E*[*v*_*t*_(*x, α*)_*i*_*v*_*t*_(*x, α*)_*j*_] − *E*[*v*_*t*_(*x, α*)_*i*_]*E*[*v*_*t*_(*x, α*)_*j*_] is going to stand for the covariance between the i-th and the j-th component of a trajectory starting in *x* and whose definitory parameter vector is *α*. Of course, the behavior of a single trajectory need not to be representative. Thus, if we are interested in studying the behavior of the model in some bounded open region of the phase space, we can define

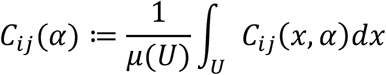

to be the covariance of the neural model specified by the parameter vector *α* averaged over the given domain, *U*. Here, it was used that *μ*(*U*) ≔ ∫_*U*_ 1*dx* is the Lebesgue measure of the open region *U*.

Finally, in order to find the correlation matrix, P, one could normalize the covariance matrix using the formula

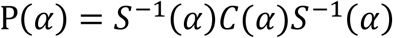

Where *S*(*α*) is the diagonal matrix whose entries consist on the variances of each component of the solution vector, this is, *S*_*ij*_(*α*) = *C*_*ij*_(*α*)*δ*_*ij*_, where *δ*_*ij*_is the Kronecker delta. From here, it could be seen that the matrix P(*α*) is defined by a continuous map of the form P: ℝ^*p*^ → ℝ^*N*×*N*^.

Thus, in order to fully characterize the matrix P(*α*) we must specify the *p* independent parameters which completely define the flow of our neural models.

In a general frame, it could be assumed that three different classes of parameters exist. In the first place, we could find parameters which affect the whole network, like the activation of a neuron of a nearby region that projects outputs to the system. These parameters do not depend on the size of the network, and thus remain constant with *N*.

Then, there are the ones that depend on individual neurons, like spike thresholds, time constants or the afferent synaptic strengths coming from outside the network. The number of such scalars increases linearly with the network size.

Finally, we have the parameters that define pairwise relations between units, like synaptic strengths between neurons of the network itself.

In general, if no specific structure is imposed on these last types of arrays, or if their components are assumed to be mutually independent, then their number grows quadratically as the network size increases. In this scenario *p*∼*O*(*N*^2^).

However, if we assume these parameters are mutually dependent, as in the case we impose connection matrices to be restricted to have a given lower dimensional rank, then their size also grows in a linear fashion, as it was shown in section 2.3, and thus one has that *p*∼*O*(*N*). Indeed, it has been already shown how the number of parameters needed to fully specify a *N* × *N* rank-*n* matrix is given by *p* = *n*(2*N* − *n*). Thus, if we also take into consideration the other types of parameters, whose cardinality cannot increase faster than linear, one could divide by *N* to find that *p*∼*O*(*N*), as claimed.

In section 2.3 it was also assured that if the dynamics were confined to a lower dimensional Euclidean manifold the same would hold true. In this case, if it is assumed the manifold is *n*-dimensional, then a PCA would find that the covariance matrix would only have *n* non-null eigenvalues, which in turn would imply that *rank*(P) = *rank*(*C*) = *n*. Then, since we already know that the complexity of a rank-*n* matrix is given by *p* = *n*(2*N* − *n*), we would also have that *p*∼*O*(*N*), as before.

